# The flow of reward information through neuronal ensembles in the accumbens

**DOI:** 10.1101/2024.02.15.580379

**Authors:** Benjamin Arroyo, Enrique Hernandez-Lemus, Ranier Gutierrez

## Abstract

Reward information flows through neuronal ensembles in the nucleus accumbens shell (NAcSh), influencing decision-making. We investigated this phenomenon by training rats in a self-guided probabilistic choice task while recording single-unit activity in the NAcSh. We found that rats dynamically adapted their choices based on an internal representation of reward likelihood. Neuronal ensembles in the NAcSh act as dynamic modules to process different aspects of reward-guided behavior. Ensembles dynamically change composition and functional connections throughout reinforcement learning. The NAcSh forms a highly connected network with a heavy-tailed distribution and neuronal hubs, facilitating efficient reward information flow. Reward delivery evokes higher mutual information between ensembles and unifies network activity, while omission leads to less synchronization. Our recordings shed light on how reward information propagates through dynamically changing ensembles of neurons in the NAcSh. These functional ensembles exhibit flexible membership, dropping in and out and even shrinking in number as the rat learns to obtain (energy) rewards in an ever-changing environment.

**Graphical Abstract:** 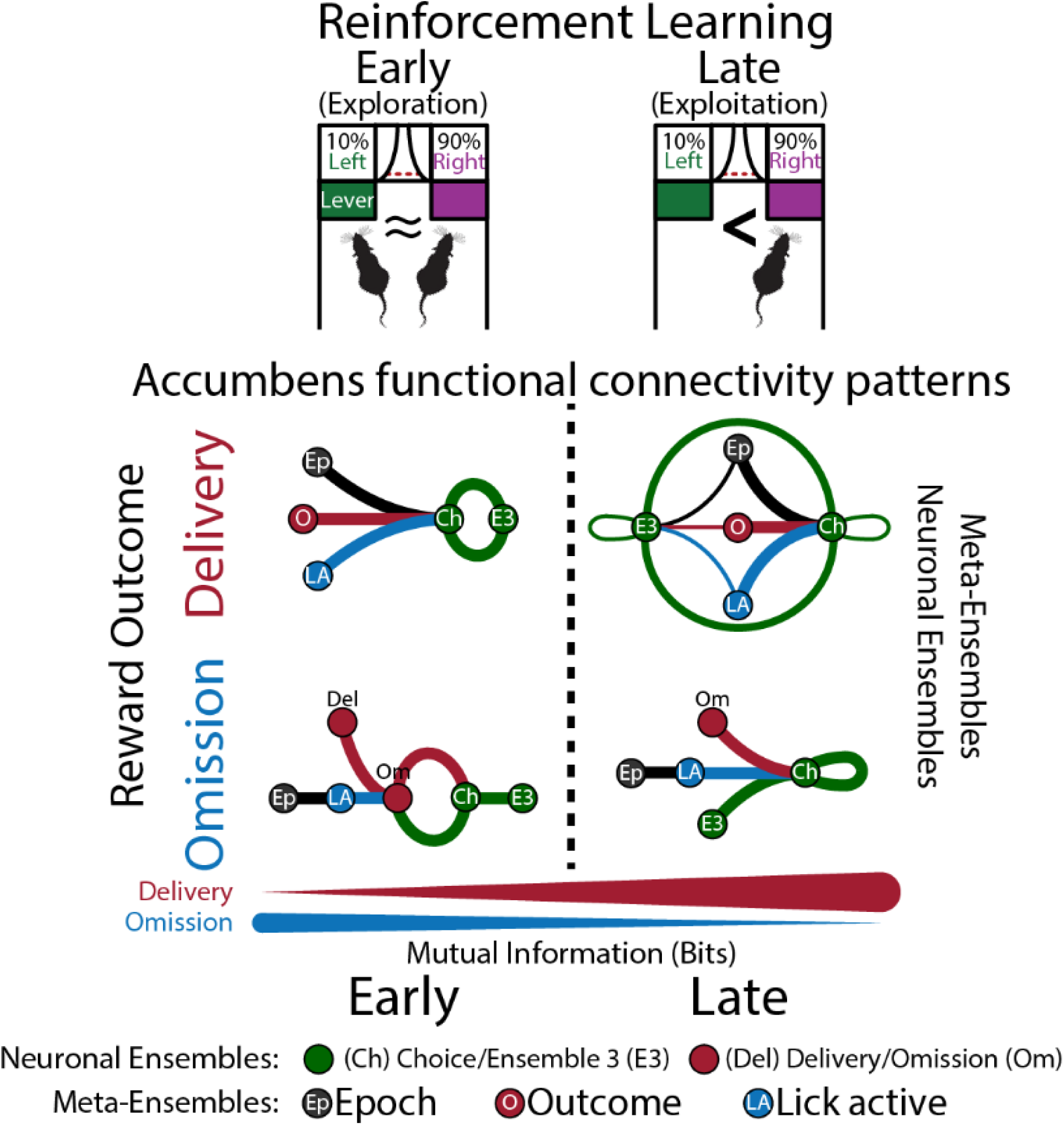

## Introduction

The nucleus accumbens shell (NAcSh) plays a critical role in reward^1,2^ and hedonic feeding.^3,4,5^ It acts as a central hub, integrating reward expectation, motivation, and decision-making signals.^1,6,7,8^ NAcSh neurons not only encode the hedonic value of rewards (sucrose palatability)^9–13^ but also guide behavior by detecting reward outcomes.^7^ Interestingly, NAcSh activity is inhibited during feeding itself,^11,14,15,16,17^ suggesting a gating mechanism that controls consumption.^5,18,19^ However, how the NAcSh separates rewarding (hedonic) signals from the oromotor components of licking is poorly understood.

The NAcSh exhibits dynamic activity patterns that are context-dependent. In simpler tasks, like freely licking for *ad libitum* available sucrose, NAcSh neurons organize into two main functional ensembles: a large, inhibited group and a smaller, activated group during consumption.^9,15^ However, for more complex tasks with limited access to tastants, like brief access tests, the NAcSh is organized into four functional ensembles that track the most relevant stimuli in the task.^9,11^ This suggests that NAc ensembles, the building blocks of neural circuits,^20–22^ increase in number and complexity with task difficulty, playing a dynamic role in decision-making by monitoring key task events. We hypothesize that this trend continues in more complex tasks like self-guided probabilistic choice. Here, animals learn reward probabilities for levers, choosing the most likely to yield a reward.^23^ This requires tracking past choices, outcomes, and the environment’s current reward state. Importantly, no external cues are guiding their actions. Instead, they must build their internal representations of reward probabilities through self-directed learning. This makes the self-guided probabilistic choice task arguably more challenging than those with external cues,^23,24,25^ demanding greater engagement in reinforcement learning and more vigilant decision-making.

This study investigates how NAcSh functional ensembles represent information in a particularly challenging task: a self-guided probabilistic choice task.^25^ Unlike tasks with external cues, this task relies solely on reward outcomes as feedback for decision-making. To understand how NAcSh encodes and processes information in this complex behavioral task, we recorded the activity of NAcSh neurons in rats performing the self-guided probabilistic choice task. This task design separates reward representation from oromotor actions (licking), allowing us to understand how reward expectation guides decision-making during reinforcement learning. We employed a multi-level analysis approach, examining the activity of single neurons, population activity, neuronal ensembles, and network activity. These analyses included techniques like time-warping, ROC analysis, PCA, t-SNE, hierarchical clustering, and network modeling to reveal the functional connections (statistical dependencies) between NAcSh neurons.

### Results

#### Behavioral results

##### Rats dynamically adjust their choices in response to reward probability and reward history

Seventeen rats were trained in a self-guided probabilistic choice task similar to the two-armed bandit reinforcement learning paradigm.^23,25^ **Figure 1A** illustrates the task structure: Each trial began with house lights illuminating, prompting the rat to visit the center port and break a photobeam with its nose. After a brief delay (250 ms), a sound (Go signal) played, and both levers extended into the box. The rat could choose either lever; importantly, the sound provided no clue which lever offered a higher reward probability.

**Figure 1.**
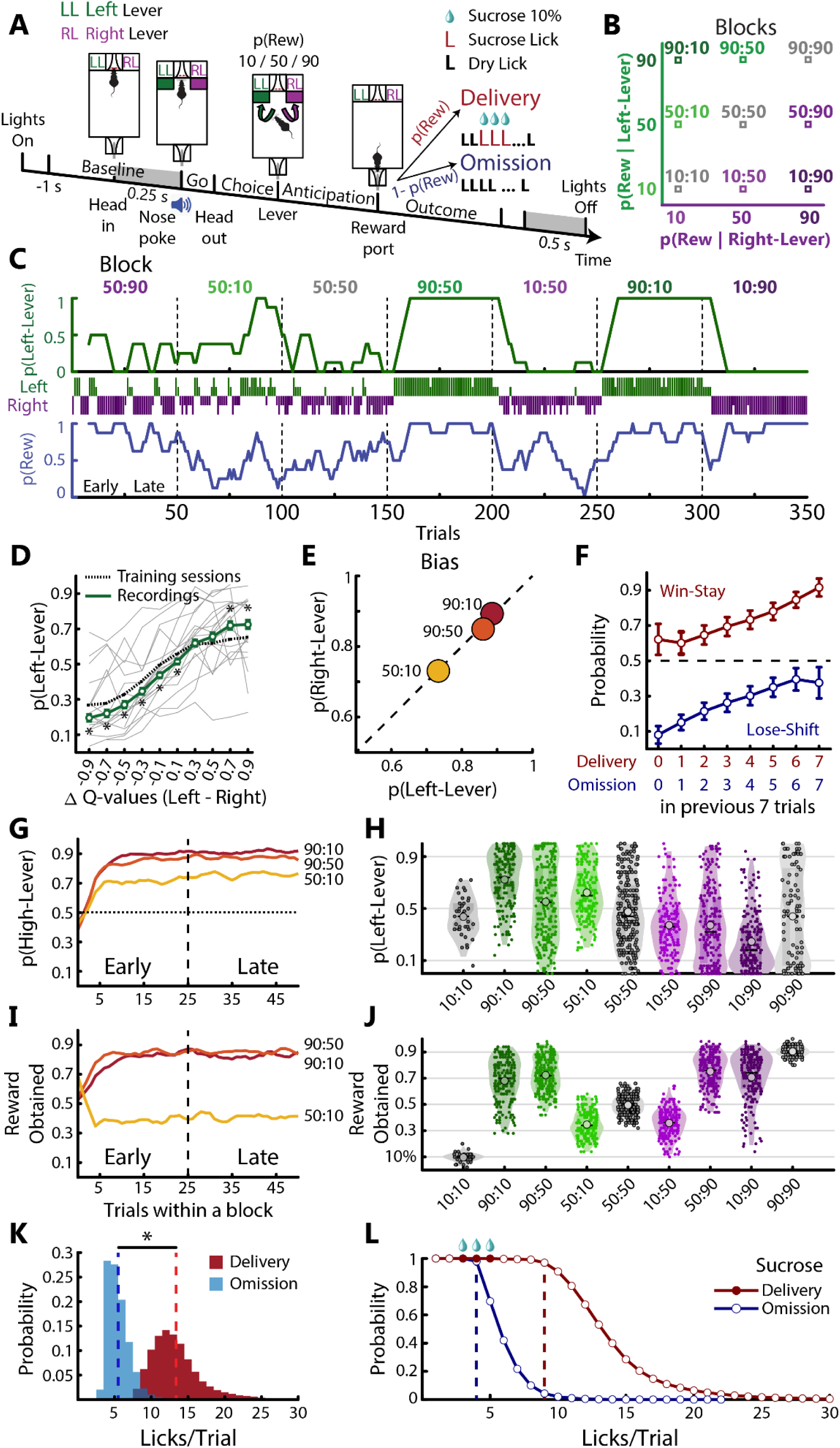
Performance of a self-guided probabilistic choice task. **(A)** A single trial structure of self-guided probabilistic choice task (see Methods). **(B)** Diagram of all reward probabilities tested. For instance, in block 90:10, 90% indicates the reward probability for the left and 10% for the right (see squares). Blocks with higher reward probability for the left lever are represented in green, purple for the right lever, and gray for blocks where the reward probabilities are equal for both levers. **(C)** Choice behavior within a representative session. Top panel, the green line, depicts the probability of pressing the left lever (smoothed over 8 trials window). In the middle panel, green and purple vertical ticks denote the left and right lever choices. Long ticks indicate rewarded trials, while short omissions. Bottom panel, the blue line depicts the reward rate. We designated the early phase as the first 25 trials of the block and the late phase as the last 25 trials. The dashed vertical lines indicate block transitions. **(D)** Left-choice probability as a function of action-value disparity. Action values were computed using the Q-learning model. Q-values near 1 indicate that the rat is more likely to choose that lever. Hence, the left lever is more likely to be pressed if the difference between Q-values (Q-value Left -Q-value Right) is positive. (*p<0.05 unpaired t-test recording vs. training, solid-green and dashed-black sigmoid, respectively). Individual rats are shown in gray lines. **(E)** Comparison between leftward and rightward blocks. No biases were found in the asymmetric blocks. **(F)** Given that no sensory cue could predict the outcome, it forces animals to integrate multiple previous trials to guide choice and promote a win-stay and lose-shift strategy^25^ **(G)** The probability of choosing the lever associated with a higher reward probability as a function of trials within a block. The horizontal dotted line denotes chance level (0.5), while the vertical dashed line signifies the transition from the Early to Late phase within each block. **(H)** Probability of choosing the left lever throughout blocks. Each small dot depicts a single session block; shaded areas are the kernel density function. Horizontal lines and big dots within violin plots indicate the median and the mean, respectively. **(I)** Reward obtained on the high-reward lever through block. **(J)** The overall rewards obtained through blocks. **(K)** Distribution of licking behavior in the reward port during sucrose delivery and omission (Delivery: 13.39 ± 0.02, Omission: 5.5 ± 0.01, red and blue vertical dashed lines, respectively, mean ± SEM). **(L)** Lick probability during sucrose delivery and omission.

The reward probabilities for the left and right levers varied across blocks (Combinations: 10%, 50%, or 90% **Figure 1B**). These probabilities switched every 50 trials with no explicit cues or sensory feedback. After choosing a lever, the rat approached the reward port on the opposite side and licked the spout twice (dry licks). On successful trials (reward delivered), the following three licks dispensed a sucrose drop (one drop per lick). Conversely, unsuccessful trials offered no sucrose (reward omission), and the rat had to lick twice more (four total). This ensured similar lick movements, regardless of the reward outcome. Importantly, the trial ended 0.5 seconds after the last lick, irrespective of the outcome. This design meant the rats, lacking external cues or warnings about changing reward probabilities, relied solely on the presence or absence of sucrose to guide their choices. In our task, we found that rats exhibited dynamic and adaptive behavior over a single session, which is the ability to change one’s behavior in response to new information or changes in the environment. Rats were more likely to choose the lever that had previously rewarded them. The more likely a lever was to give a reward, the more likely the rats chose it (**Figure 1C**).

To model the rat’s choice probability, we used Q-learning, a value-based reinforcement learning model. This approach uses the subject-specific history of rewards and actions to estimate action values, commonly called Q-values. In the rat’s current state, the Q-value associated with an action reflects the expected value of taking that action. The increased Q-value of the left lever correlates with the higher likelihood of the rat choosing that option. The relationship between the choice probability and the delta of Q-values further improved after electrode implantation (as indicated by comparing the dashed black sigmoid curve with the solid green sigmoid curve in **Figure 1D**). Thus, electrode implantation did not impair the task’s performance; rats kept improving. In **Figure 1E**, there was no significant preference (bias) towards either the left or right lever selection in the blocks highly rewarded leftward or rightward (Unpaired t-test, 50:10 vs. 10:50 t _(246)_ = 0.44, p =0.66; 90:50 vs. 50:90, t _(171)_ = 0.54, p =0.54; 90:10 vs. 10:90, t _(162)_ = −0.5, p =0.61). This result implies that rats did not inherently prefer one lever over the other, suggesting that the likelihood of receiving a reward was mainly responsible for guiding the rat’s decisions.

To understand how reward delivery or omission influenced the rats’ choices, we examined how reward history affected their decisions. The results showed a clear pattern: as the number of times a lever yielded a reward increased; the rats were more likely to pick that same lever again (Kruskal-Wallis test; stay probability: χ^2^ _(7, 1985)_ = 636.43, p<0.0001; **Figure 1F red**). This tendency is called the win-stay strategy. Conversely, when a lever provided fewer rewards, the rats became more likely to switch to the other lever (Kruskal-Wallis test, shift probability: χ^2^ _(7, 1985)_ = 6.36.98; p<0.0001; **Figure 1F** blue). This is known as the lose-shift strategy. These findings suggest the rats were sensitive to reward history and used a win-stay, lose-shift strategy to guide their choices.

The reward probabilities changed between blocks without warning, forcing the rat to update its beliefs about the current reward probability state in the new block.^26^ This triggered an initial exploration phase, in which the rat focused on gathering information. They likely relied on previous trials to assess the potential value of each lever (Early phase). After gathering enough information, the rats transitioned to an exploitation phase. Here, they aimed to maximize rewards by choosing the lever with a higher probability of success. This latter part of the block was designated as the Late phase.

As anticipated, rats initially showed a reduced preference for the higher-rewarding lever during the Early phase of each block. However, as they gained experience within the block, their preference for this lever increased significantly in the Late phase. This effect was most pronounced in blocks with a higher potential for greater overall reward (90:50 and 90:10; **Figure 1G**). It follows that the exploitation of learned action values (post-learning) in the Late phase may be less cognitively demanding than the exploration behavior in the Early phase that supports new learning.

Figure 1H shows the probability of choosing the left lever across blocks. Note that rats chose the left lever more often in blocks where it had the higher reward probability (green) and chose the left lever less often in blocks where the higher reward was assigned to the right lever (purple; Kruskal-Wallis test, p(Left-Lever): χ^2^ _(8, 1516)_ =361.14, p<0.0001). In blocks with equal reward probability (gray blocks), rats exhibited a bias, responding more to one lever than the other. However, across subjects, this effect canceled each other out. Interestingly, in the block with the lowest reward rate (Figure 1H, see 10:10), rats were more likely to explore and constantly visit both levers than in the 50:50 and 90:90 blocks.

We also noted a proportional increase in rewards obtained to the highest reward probability in each block (Figure 1I). This observation was accurate not only for blocks highly rewarded leftward or rightward but also across the other blocks (Figure 1J; Kruskal-Wallis test, Reward rate: χ^2^ _(8, 1516)_ =1034.88, p<0.0001). Our findings indicate that rats could detect and adapt their choices in response to an environment characterized by nonstationary reward probabilities.

We investigated how reward delivery or omission affected licking behavior in rats. Rats licked significantly more during rewarded trials compared to unrewarded trials (Unpaired t-test, t _(79899)_ = 347.5; p <0.001; Figure 1K). In 97% of rewarded trials (n = 47,007), rats licked at least nine times (Figure 1L dashed red line). In contrast, in 98% of the trials where the reward was omitted (n = 32,894), they emitted (as requested) at least four dry licks (see the dashed blue vertical line in Figure 1L). This rapid cessation of licking after four attempts in unrewarded trials suggested that rats can detect the absence of reward within a single lick.^27^ This highlights their remarkable ability to learn and adapt to changes in reward probability. Essentially, rats track their choices, demonstrate high sensitivity to reward outcomes, and adjust their behavior based on updated expectations, all based solely on the presence or absence of reward.

### Electrophysiology

#### Single neuron analysis

##### Individual neurons encode choice, reward outcome, and rhythmic licking

A total of 1507 single neurons were recorded in the anterior NAc shell (NAcSh; **Figure S1**) while rats performed a self-guided probabilistic choice task. A time-warping algorithm realigned the neural activity across trials to account for variations between time events induced by the rats’ pace.^28^ This technique essentially stretched or compressed the time between specific task epochs (**Baseline**, **Go**, **Choice**, **Anticipation**, **Outcome**) to match the median duration observed across all trials (see **Figure S2**). By minimizing time lag variability, this preprocessing step allowed us to analyze neural activity throughout the entire trial structure (**Figure S3**).

We used receiver operating characteristic (ROC) curves to investigate if individual neurons fired differently based on the choice (left vs. right lever) or outcome (rewarded vs. unrewarded) during the task. Our analysis revealed significant responses in NAcSh neurons linked to these variables. For example, one subset of neurons increased their firing rate as the rat approached the reward port, but this activity was inhibited just before licking. This neuron then exhibited a phasic activity during reward delivery but decreased activity when no reward was given, suggesting they encode reward outcomes (Figure 2A, bottom panel, red arrow). We named them “Outcome neurons.” Similarly, we found neurons that responded specifically to the choice of lever (left vs. right). Some neurons could even encode choice and outcome (Figure 2B). The top panel of the Peri-Stimulus Time Histogram (PSTH) shows modulation by choice (green arrow), while the bottom panel indicates response to reward outcome (red arrow). These neurons were termed “Choice & Outcome neurons.” Finally, we identified neurons whose activity covary with licking behavior (Figure 2C), called “lick-coherent neurons.” Neurons that encoded all three variables (choice, outcome, and licking) were less frequent (more of this below).

**Figure 2.**
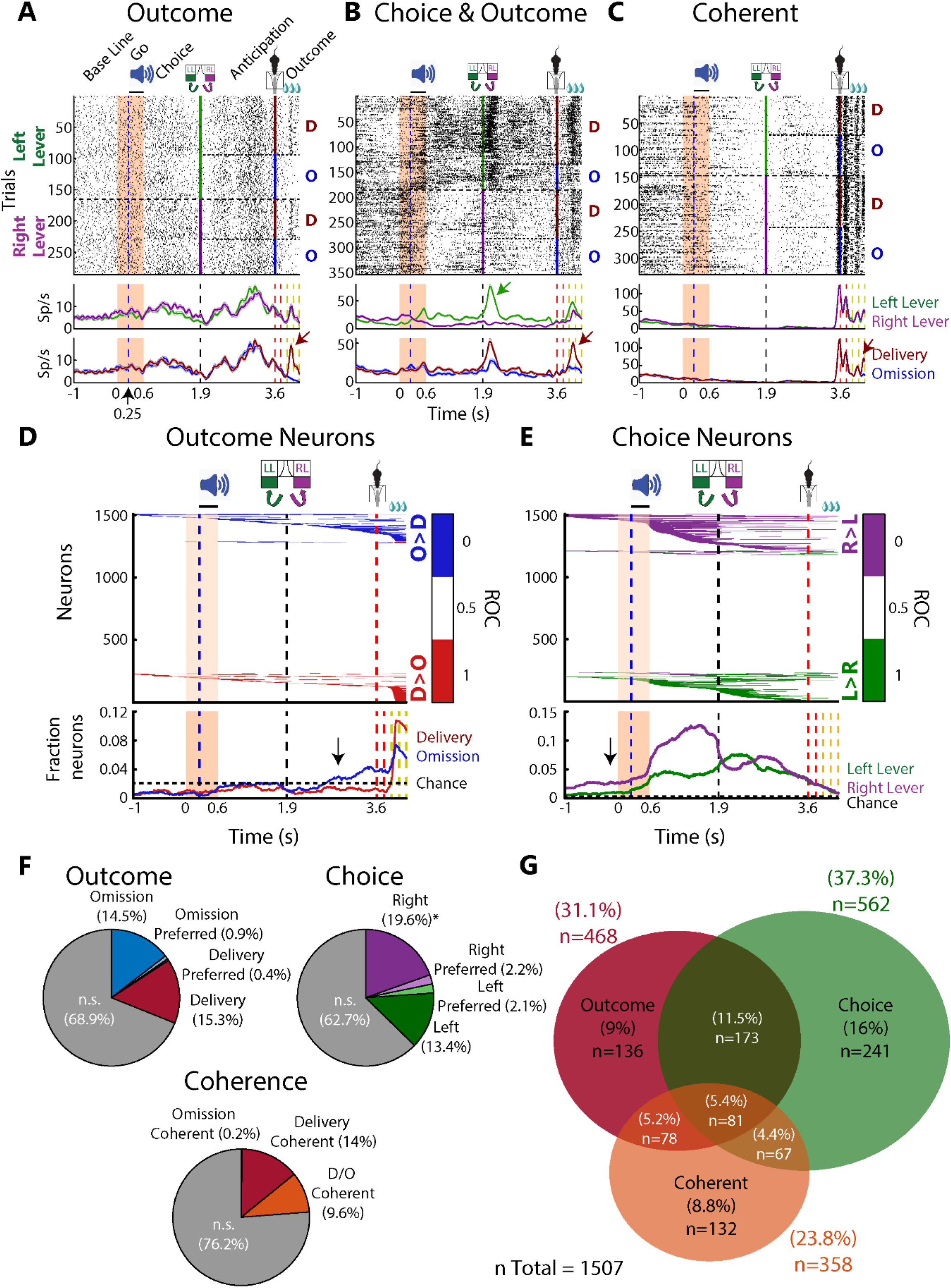
Extracellular single-unit recordings identified NAcSh responses that tracked the choice, reward outcome, and rhythmic licking. **(A)** Single neuron responses using a time warping allow us to visualize the activity across epochs (**Figure S2**)^28^. The raster plot depicts the time-warped response of an Outcome neuron that fired more for sucrose delivery compared to omission (see red arrow). The raster plot is aligned to the head entry (Time = 0 s). The blue dashed line marked the “Go” tone (Time = 0.25 s), and the head out is shown at time = 0.6 s. Lever presses are displayed at 1.9 s. Trials were sorted by left or right lever press (green and purple vertical lines, respectively), which were sorted by reward delivery (D; red ticks) and omission (O; blue ticks). The 1^st^ lick delivered in the reward port is shown at 3.6 s: The middle panel indicates the PSTH of neuronal activity for left and right lever press trials (green and purple lines, respectively). The bottom panel is the PSTH responses for reward delivery and omission (red and blue lines, respectively). Red vertical dashed lines indicate the two empty licks and yellow dashed lines represent sucrose drops for delivery trials (one per lick), whereas, in omission, they are empty. **(B)** A representative NAcSh neuron is selectively activated for the left lever press (green arrow) and sucrose delivery (red arrow). Such responses were classified as choice & outcome neurons. Same convention as panel (**A**) **(C)** Coherent firing of a representative neuron entrained by licking behavior. This lick-coherent neuron exhibited oscillatory activity that decreased amplitude as the licks were given in the reward port (red arrow). **(D)** Outcome neurons identified by the ROC curve. Top panel, blue bins indicate the time intervals where neurons fired significantly more during reward omission trials compared to delivery trials (O>D), while red bins represent significant time intervals with greater activity for reward delivery trials (D>O). Significant responses were sorted by onset latency. The bottom panel, red, and blue lines, represent the fraction of selective neurons for reward delivery and omission, respectively. The horizontal dashed line indicates the chance threshold calculated by permutations (see Methods). **(E)** ROC analysis for Choice neurons. Top panel, purple bins represent time intervals for neurons selectively active for trials where the right lever was pressed (R>L), while green bins indicate time intervals for trials where the left lever was pressed (L>R). Bottom panel depicts the fraction of neurons recruited for left (green line) and right (purple line) lever presses over time. **(F)** Pie charts depicting the percentage of neurons modulated by Outcome, Choice, and Coherence. **(G)** Veen’s diagram illustrates the logical relations between Outcome, Choice, and Coherent-related neuron sets.

Our findings revealed that most neurons in the NAcSh exhibited robust responses selective to reward outcomes, particularly during the Outcome epoch. We also observed neurons that fired selectively in anticipation of omitted rewards (Anticipation epoch, blue line, black arrow in Figure 2D). This is noteworthy because there were no cues signaling the omission of the reward. This implies a potential role of these neurons in expecting and predicting the omission of reward.

Similarly, NAcSh neurons exhibited responses related to the choice, spanning the Choice and Anticipatory epochs. A subpopulation of choice-selective neurons anticipated the rat’s decision regarding which lever to press. Most of these neurons were recruited after the Go signal that preceded the lever press. Interestingly, a few neurons carried choice-related information even during the Baseline epoch (i.e., before the trial began, indicated by a black arrow in Figure 2E). We propose that these anticipatory responses might reflect the subjects’ probability of making a forthcoming choice.^29^

Outcomes modulated 31.1% of NAcSh neurons (468/1507). Specifically, 15.3% of NAcSh neurons (n=230) were selectively activated during reward delivery, while 14.5% (n=219) responded only to reward omission (Figure 2F, Delivery and Omission). Neurons modulated by both reward delivery and omission at different time intervals were classified as either “Delivery preferred” (more significant bins for delivery) or “Omission preferred” (more significant bins for omission). Thus, we identified 0.4% of neurons (n=6) as Delivery preferred, while 0.9% (n=13) were classified as Omission preferred (Figure 2F, Outcome pie chart).

Choice encoding was significant in 37.3% of recorded neurons (562/1507). Within the NAcSh population, a significant bias towards right-lever selectivity was observed, with 19.6% (n=295) of neurons showing activation for the right-lever choice, compared to only 13.4% (n=202) for the left-lever choice (χ^2^=20.39, p<0.0001). The observed bias towards right-selective neurons may be because of the larger sample size from the right hemisphere (1186 vs. 321 neurons). This is supported by the exclusive presence of a significantly greater proportion of right-selective neurons in the right hemisphere (**Figure S4**). We found neurons significantly activated by left and right choices but at different time intervals, only 2.1%. of NAcSh neurons (n=32) were left-preferred, and 2.2% (n=33) were right-preferred (Figure 2F, Choice pie chart).

Regarding the lick-coherent neurons, 23.8% of neurons (358/1507) exhibited significant coherence with rhythmic licking. Most of these neurons were lick-coherent either only in delivery trials (14%, n=211) or in both delivery and omission trials (D/O Coherent; 9.6%, n=144), while only a few neurons (0.2%, n=3) were coherent only in omission trials (Figure 2F, see Coherence pie chart).

We observed that NAcSh neurons were selective to one or more variables, with major overlap found in neurons modulated by outcome and choice (11.5%, n=173), followed by a small subpopulation modulated by all three variables (5.4%, n=81; Figure 2G). Conversely, a subset of lick-coherent neurons also conveyed information related to Outcome and Choice. For instance, 5.2% of neurons (n=78) were modulated by outcome and licking behavior, while 4.4% (n=67) were modulated by choice and licking behavior (**Figure S3C**). This result suggests that these lick-coherent neurons were more complex than those that conveyed pure oromotor responses (**Figure S3C**). Finally, only 7.8% of neurons (n=119) were not responsive to any of these variables. These findings indicate that most NAcSh neurons (92.2%, n=1388) were engaged in the task, multiplexing two or more variables. In summary, the recruitment of outcome, choice, and lick-coherent neurons provides valuable insights into how single neurons in the NAcSh encode consummatory and reward-related information to guide decision-making.

#### Population analysis

Principal Component Analysis (PCA) was employed to uncover the neuronal trajectories within NAcSh population activity in response to reward outcomes. This analysis revealed that distinct patterns associated with the delivery and omission of rewards were reflected in a linear low-dimensional state space of neuronal population activity.

##### NAcSh activity diverged into two distinct neural state spaces, reflecting both outcomes

Using PCA,^30^ we investigated how populations of neurons in the NAcSh collectively encode behavior. While the first principal component (PC1) related to licking behavior (29.89% of variance, Figure 3A) did not differentiate between reward outcomes, suggesting NAcSh is more related to oromotor responses. PC2 mirrored PC1, albeit negatively, further supporting its association with licking. In contrast, PC3 and PC4 trajectories distinguished between outcomes: PC3 neurons were most active during reward delivery (Figure 3A, red), while PC4 favored reward omission (blue). This suggests that processing reward outcomes are the second most prominent variable tracked by NAcSh at the population level.

**Figure 3.**
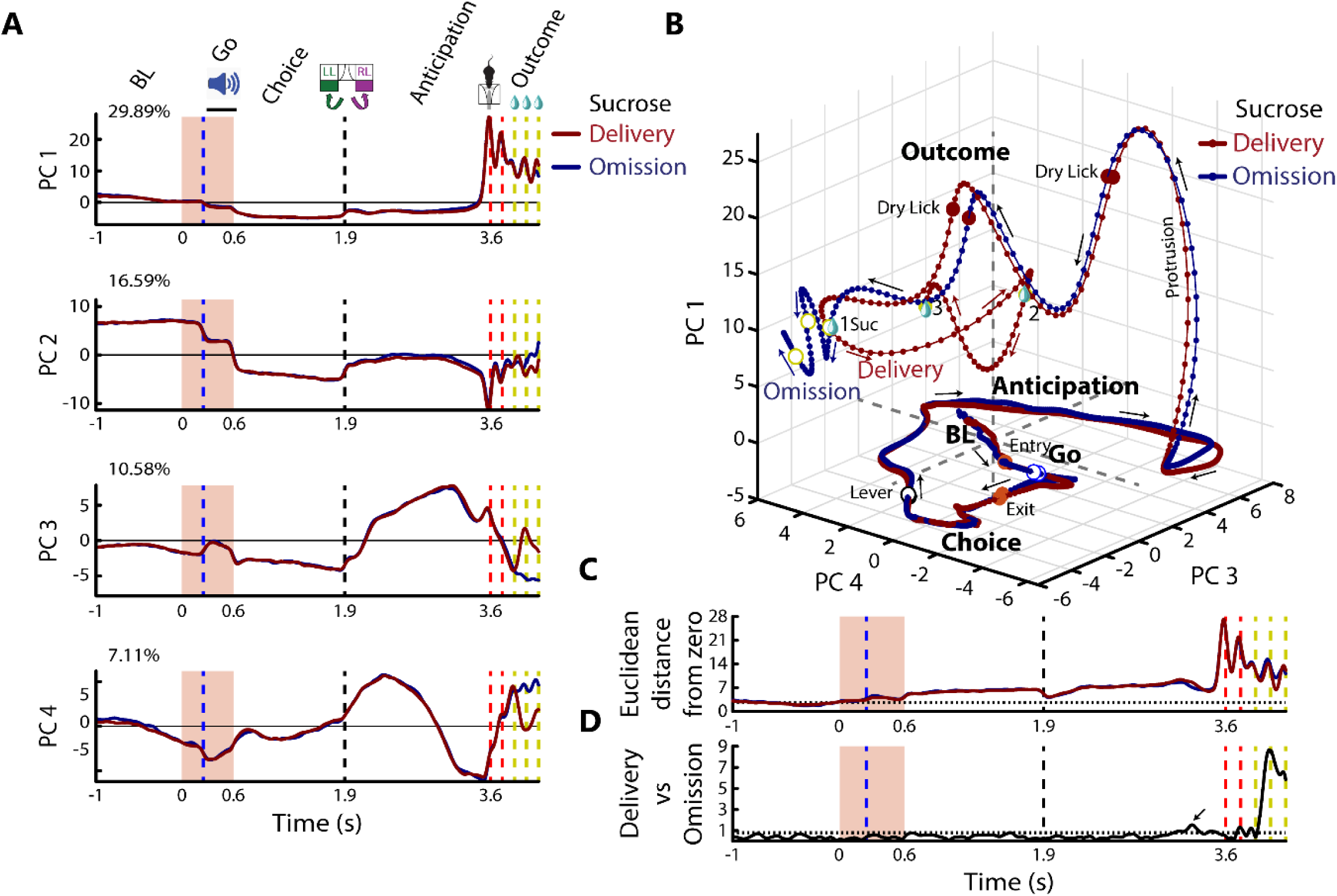
PCA trajectories of the NAcSh discriminated between reward outcomes. **(A)** PCA of neuronal population dynamics across task epochs during reward delivery (red) and omission (blue). The percentage of the variance explained by each component is also shown. **(B)** The PCA trajectories for delivery and omission trials (red and blue dotted lines, respectively). The arrows indicate the direction in which trajectories followed across epochs. **Video S1** details PCA trajectories as a function of neuronal ensembles (identified in Figure 4). **(C)** Euclidean distance for reward delivery and omission trajectories (red and blue lines, respectively). The horizontal dotted line represents the mean distance value during the baseline epoch. **(D)** The black line represents the Euclidean distance between outcome NAcSh PCA trajectories. The separation of reward vs. oromotor effects occurs precisely at reward/omission, with an anticipatory effect noted (arrow). The dotted line indicates the maximum baseline distance

NAcSh activity tracks reward processing in three PCA dimensions (PC1, PC3, PC4), explaining nearly half (47.58%) of the total variation in firing rates across neurons. Figure 3B reveals three PCAs that effectively capture the dynamic activity of neurons in the NAcSh during reward anticipation, licking, and outcome processing. It is important to note that until the outcome epoch, the NAcSh population activity followed similar paths throughout the trial task (Figure 3B). However, within the outcome epoch, a striking shift emerges. Here, the rhythmic licking of the empty sipper becomes tightly coupled to oscillations in the population activity.^31^ Interestingly, these oscillations diverge into distinct state spaces for rewarded and unrewarded trials, effectively separating reward processing from the act of licking.

To quantify the overall modulation level, we calculated the Euclidean distance between each time bin and the origin of the PCA space (Figure 3C). This distance reflects the overall “perturbation” of the population activity from its baseline state. As illustrated in Figure 3C, the PCA trajectories progressively move away from the origin (stable state) after the Go signal. Positive values on the y-axis indicate a more substantial perturbation, with distance steadily increasing through the Choice epoch. Within the Anticipation epoch, the Euclidean distance peaks just before the first empty lick (tongue protrusion) in both reward outcomes. This highlights the progressive recruitment of neurons as the animal expects and attempts to receive a reward, regardless of the actual outcome.

In the outcome epoch, PCA trajectories oscillate in sync with dry licks (until the 3^rd^ lick). However, an important shift occurs when a reward is delivered (1^st^ rewarded lick). As noted, distinct neural state spaces emerge, separating rewarded and unrewarded trials (Figure 3B). To measure this divergence, we calculated the Euclidean distance but now between rewarded and unrewarded trajectories (Figure 3D). Initially, they follow similar paths (across epochs), but a slight divergence during the anticipation epoch hints at reward anticipation (see arrow). This divergence sharply increases after the first reward delivery (yellow dashed line), highlighting the separation of licking-related activity from the reward response itself (**Video S1**). Our population analyses suggest that the NAcSh monitors trial progression and distinguishes reward outcomes, highlighting the NAcSh’s key role in encoding the anticipation and the outcome of actions.

##### Neuronal activity patterns in the NAcSh also encode reward history and choice strategy

Since our task does not provide external cues indicating the subsequent reward outcome, rats cannot be certain if a reward will be omitted on a single trial. However, rats could generate a reward expectation based on reward history (see Behavior Figure 1F; win-stay & lose-shift). This expectation may influence their choice strategy. To investigate whether neuronal activity encodes reward history, we performed a PCA analysis considering reward history (last three trials) and the rat’s choice strategy (stay vs. shift). Our analysis revealed that the NAcSh population trajectories also encoded reward history, choice strategy, and their interaction (see **Figure S5**). Population trajectories distinguished reward history and choice strategy throughout the trial, from baseline to anticipatory epochs, but rapidly converged during the outcome epoch.

#### Ensemble analysis

##### The heterogeneity of NAcSh responses is organized into “functional” neuronal ensembles and meta-ensembles

Leveraging PCA, t-SNE, and hierarchical clustering, we identified 14 distinct neuronal ensembles (initially named A1-A14) in the NAcSh (Figure 4A, see the Ensemble outcome map, the letter “A” indicates that “All” 50 trials in the block were used to compute t-SNE). When we identified a plausible “function” of the neuronal ensemble, we renamed it accordingly. These 14 ensembles were automatically determined using the elbow method (see Methods). These ensembles were functionally categorized into four meta-ensembles through a manual process (color-coded contours in Figure 4A) based on their location on the t-SNE map, firing patterns, and relation to task events:

**Figure 4.**
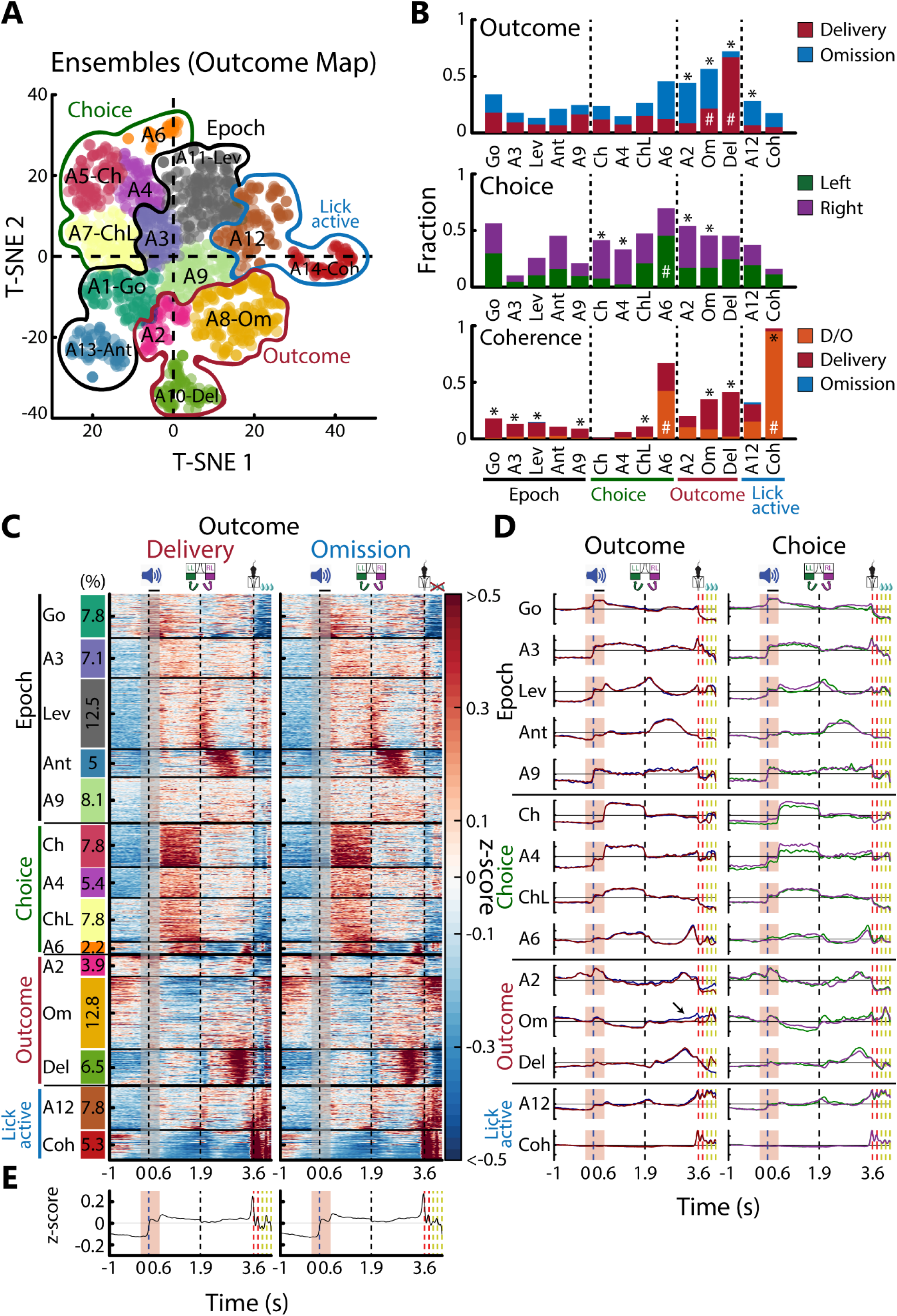
The diverse activity patterns of NAcSh responses were grouped into 14 neuronal ensembles, each specializing in processing specific task- and reward-related information. **(A)** Outcome ensemble map was generated using t-distributed stochastic neighbor embedding (t-SNE) of time-warped firing rates (see Methods). Each dot represents a single neuron, color-coded according to the ensemble. Meta-ensembles are also depicted: Epoch (black contour), Choice (green), Outcome (red), and Lick Active (blue). **(B)** Bar plot illustrating the fraction of neurons encoding Outcome (top, using ROC), Choice (middle), and Coherence (bottom) variables across ensembles. Ensembles are sorted based on neural activity patterns, forming different Meta-ensembles. The hash symbol denotes ensembles with a larger fraction of neurons selective to a variable (# χ^2^ test, p<0.05), while the asterisk symbol (*χ^2^ test, p<0.05) indicates statistically significant differences between conditions (e.g., delivery vs. omission, left vs. right). **(C)** Colored-coded heatmaps for neurons assigned to fourteen ensembles. Each row represents the z-scored mean activity from a single neuron during reward delivery (left) and omission trials (right). Neuronal ensembles are sorted as in **B**, and the vertical bar color on the left indicates the percentage of neurons for each neuronal ensemble. All responses were aligned to the head entry (onset of gray shaded area). From left to right, the first vertical dashed line is the Go signal, the second the lever press, and the third is the first lick delivered in the reward port. Solid, thick horizontal lines separate individual ensembles and meta-ensembles, respectively. **(D)** Average ensemble activity with distinct neuronal activity patterns identified by t-SNE and sorted by Outcome and Choice (left and right panels, respectively). Neuronal ensembles include Go (A1-Go), A3, Lev (A11-Lev), Ant (A13-Ant), A9, Choice (A5-Ch), A4, Choice-Late (A7-ChL), A6, A2, Omission (A8-Om, the arrow indicates the increased activity preceding the omission of the reward), Delivery (A10-Del), A12, and Coherent (A14-Coh). A more detailed PCA analysis of the Omission ensemble (A8-Om) revealed that it also conveys reward history information (see **Figure S6**). **(E)** The black line represents the average population activity in the NAcSh for reward delivery and omission trials (left and right panels, respectively).

The *Epoch meta-ensemble* (black contour) comprises five neuronal ensembles: A1-Go (n=117) signals the initiation of the trial, and thus, we renamed it as the Go ensemble. A3 (purple, n=107) fires more during go and choice epochs, and A11-Lev (n=189) is an ensemble responsive to lever press. A13-Ant (n=75) fires during the anticipation epoch, and the A9 (n=122) is a goal-directed ensemble because it exhibited a sustained activity across multiple epochs, from Go, Choice, and Anticipation epochs, probably monitoring the entire reward-seeking sequence. An Epoch meta-ensemble could track all-important task events and epochs.

The *Choice meta-ensemble* (green) is formed by four neuronal ensembles named A5-Ch (Choice, n=118), A4 (ensemble 4, n=81), A7-ChL (Choice Late, n=118), and A6 (ensemble 6, n=33). All these ensembles fired more during the Choice epoch, but as noted below, they all have information about choosing the left or right lever and reward omission.

*The Outcome meta-ensemble* (red) consists of three ensembles: A2 (ensemble 2, n=59), Om (Omission, n=193), and Del (Delivery, n=97). A2 and Om ensembles fired more on omission trials, while the Del ensemble was selective to reward delivery.

*The Lick-Active meta-ensemble* is composed of two neuronal ensembles specifically associated with licking behavior. A12 (ensemble 12, n=118) and Coh (lick coherent, n=80) were the only two ensembles that exhibited more activity during licking; thus, we called it Lick-Active (LA) meta-ensemble since it provides a neural representation of the active licking process.

Next, we analyzed the distribution of neurons encoding task variables (identified by significant ROC curve analysis) across different ensembles (Figure 4B). In the Epoch meta-ensemble, few neurons encoded outcome, choice, or licking variables, except for the “Go” ensemble with more choice-selective neurons. Conversely, the Choice meta-ensembles showed a higher proportion of neurons carrying choice information and lick-coherent activity, particularly ensemble A6. The Outcome meta-ensemble had more neurons associated with reward outcomes, with two distinct patterns: Ensembles A2 and Om favored reward omission, while Ensemble “Del” favored reward delivery. The Lick-Active meta-ensemble contained some outcome/choice-encoding neurons, but the “Coh” ensemble was particularly enriched in lick-coherent neurons. These findings reveal that lick-related activity is concentrated in specific ensembles, while choice and outcome information is more broadly distributed across neuronal ensembles.

Figure 4C illustrates the normalized activity of recorded neurons, organized into 14 ensembles and subsequently grouped into four meta-ensembles, using a color-coded heatmap. Trials with reward delivery (left panel) reveal the temporal dynamics of ensemble activation during reward anticipation and receipt, while trials with reward omission (right panel) highlight ensemble responses to the absence of reward. This visualization elucidates the organization of population activity into distinct neuronal ensembles exhibiting similar activity profiles (Figure 4C). Figure 4D, the left panel plots the ensemble peri-stimulus time histograms (PSTHs), segregated by outcome delivery (red) and omission (blue), revealing distinct neuronal patterns that implicate different roles in reward processing. The right panel displays ensemble activity but is now sorted by choosing the left (green) or right (purple) lever. For example, besides encoding choice, Ch, A4, and A6 ensembles (Figure 4D) also respond more to reward omission. This ensemble activity resembles individual responses of Choice&Outcome neurons (Figure 2B). In addition, the Om ensemble also exhibited anticipatory activity that predicts the absence of reward (see Om blue line and arrow Figure 4D), but during reward delivery, it fired briefly after the first drop of sucrose. As noted above, the Om ensemble’s “pessimistic” activity pattern might also reflect information about reward history. We conducted a PCA analysis solely on its activity to investigate how the Omission ensemble captured the rat’s reward history (last three trials). Our analysis revealed two components, PC1 and PC5, that exhibited increased activity as more rewards were omitted in the past. These patterns were evident throughout the trial, persisting until the first lick in the outcome epoch (see **Figure S6**). This finding suggests that the Omission ensemble predicts the absence of reward carrying information about the rat’s recent reward history. In sum, each ensemble’s unique spatiotemporal signature provides insights into its potential function. Average population activity (Figure 4E) reveals NAcSh modulation throughout the task, with an anticipatory peak before licking, followed by sucrose-induced inhibition (though still above baseline). These results highlight the diverse contributions of distinct neuronal ensembles to reward-guided behavior, with each ensemble acting as a specialized module for processing different aspects of the task.

#### Network analysis

##### Functional connectivity of the NAcSh network exhibited a small-world architecture with a heavy-tailed distribution

We employed a probabilistic graphical model, specifically an undirected network, to analyze the functional connectivity between NAcSh neurons during different task conditions and trial types. Here, a functional connection refers to significant statistical dependency between the activity of two neurons that describes how much these neurons modulate together, even if they are not necessarily physically connected via a synapse or gap junction. Functional connections between neurons were calculated using pair-wise mutual information (MI), in which the 90^th^ percentile of the distribution was the upper cutoff to consider a functional connection; otherwise, a connection was rejected (**Figure S7**). Thus, we obtained a probabilistic graphical model, which revealed the architecture of functional connections within the NAcSh during reward outcomes (Figure 5A).

**Figure 5.**
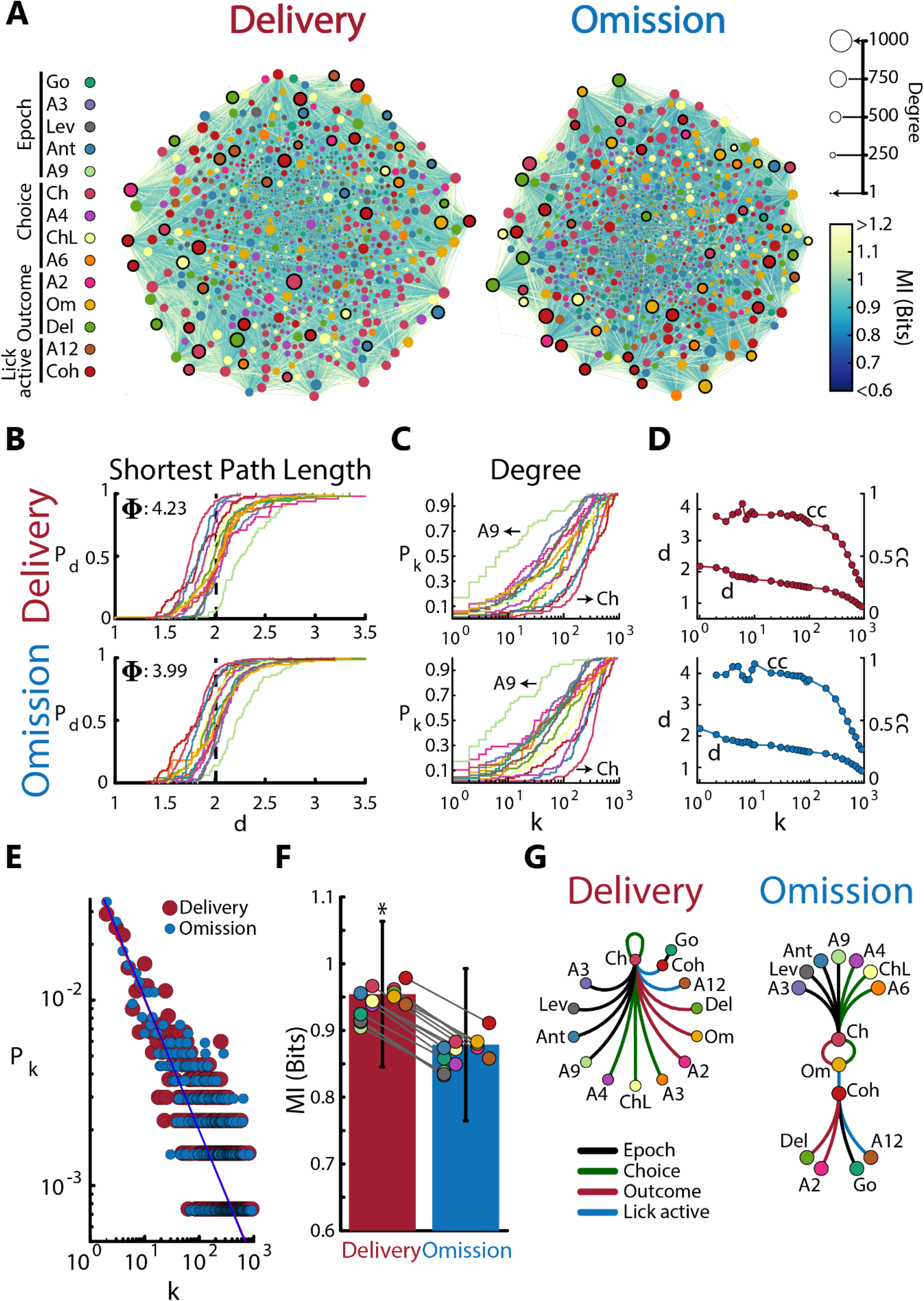
The NAcSh activity follows a small-world architecture with scale-free-like properties, and reward delivery enhances the mutual information shared across the network. **(A)** The NAcSh network during reward delivery and omission. The network is illustrated by an organic layout (computed with Cytoscape), in which each neuron is represented by a node (circle) and a ‘functional’ connection by a line between a pair of nodes. Nodes are color-coded according to the ensemble membership. The size of the node depicts the number of connections (Degree). The color of the connections indicates the mutual information shared among nodes (see **Figure S7** for the complete adjacency MI matrix). Nodes with a thick black border represent nodes with high betweenness centrality, defined as how often a neuron lies on the shortest paths between any pair of nodes in the network. **(B)** Cumulative density function of distance (d) in the NAcSh network. Lines represent distance cumulative density function across ensembles during reward delivery and omission. A vertical black dashed line indicates the mean distance in the NAcSh network. The network size was denoted by Φ. **(C)** Cumulative density function of degree connectivity in the NAcSh network. Lines depict degree cumulative density function across ensembles in reward delivery and omission (top and bottom panels, respectively). **(D)** The NAcSh network follows a small-world architecture. The distance (d, left axis) and Clustering Coefficient (cc, right axis) are both functions of degree connectivity during reward delivery and omission (red and blue dotted lines, respectively). **(E)** A heavy-tailed distribution was found in the NAcSh network. Red and blue circles depict the degree distribution for reward delivery and omission, respectively. A power-law function was fitted (Delivery γ=0.69, Omission γ=0.70; more details in **Document S1**). **(F)** Stronger Mutual Information was observed in the NAcSh network during sucrose consumption. Red and blue bars depict mean MI in the NAcSh network during reward delivery and omission, respectively. Solid black lines indicate the SEM. Color-coded circles represent the mean MI across ensembles. **(G)** Reward outcomes reorganized functional connectivity patterns. Graphical representation of the 1^st^ strongest functional connections in the NAcSh network during reward delivery and omission (more detail in **Figure S10**). Color-coded circles represent neuronal ensembles. The strongest functional connections are indicated by lines and colored according to the meta-ensemble color.

Next, we quantified some topological properties of the neuronal networks during reward outcomes. We found a similar number of neurons connected (Delivery N=1337 vs. Omission N=1358) and a similar number of connections in the Delivery and Omission networks (L=112799 vs. L=112800). We then evaluated the network properties regarding nodes. We measure the distance (d) between neurons in the network by the average shortest path length (SPL). We found that when selecting two random neurons in the network, the average distance was 2 steps, and the largest distance, or network size, was approximately 4 steps in both reward outcomes (Figure 5B). This result indicates the high interconnectivity in the network, reflecting “short-cuts” that facilitate information propagation.

We then investigated whether functional connections were evenly distributed among neuronal ensembles. We found a coexistence of many neurons with few functional connections and a few highly connected neurons, termed hubs. We also identified neuronal ensembles with a higher likelihood of having functional connections (Figure 5C, compare A9 poorly connected vs. Ch richly connected). Unlike individual neurons, neuronal ensembles appear to follow a hierarchical organization, with some ensembles being more connected than others. In particular, the Ch ensemble was consistently the most connected (see also **Figure S8**).

Furthermore, out of 1507 neurons, we identified only 57 neuronal hubs. Most hubs comprised neurons encoding outcome and choice, or more commonly, were lick coherent. However, some hubs did not encode any of these variables. Interestingly, the neuronal activity of hubs was not limited to the outcome epoch, as they exhibited various responses throughout the trial (see **Figure S9**).

Then, we were interested in how the degree of connectivity affected the distance (d) between neurons and the clustering coefficient (cc, i.e., a measure reflecting the tendency of a triad of nodes to form a complete triangle). We observed that the distance between neurons decayed rapidly as the degree of connectivity increased. In contrast, the clustering coefficient decayed only with hubs, indicating that most neurons had high local connectivity (Figure 5D). In other words, the predominant short-cuts and the higher local connectivity (high cc) in the network facilitate the information flow, indicating that the NAcSh activity follows a “small world” architecture.^32^

Once we had evidence of neuronal hubs, we evaluated whether the degree distribution followed a long-tailed distribution. The slope of the power-law function fit was similar in the NAcSh network regardless of the outcome (γ=0.69 vs. γ=0.70, red and blue lines, respectively; Figure 5E). This result seems to support a scale-free-like property in both networks.^33^ A more detailed hypothesis testing and goodness-of-fit calculation analysis can be found in **Document S1**.

##### Sucrose delivery enhances the mutual information shared across the network

Our analysis of mutual information in the Delivery and Omission network revealed a significant difference. Mutual information measures how much knowing the activity of one neuron reduces uncertainty about the activity of the other. In essence, it reveals the amount of information shared between the two neurons’ firing patterns. The mutual information in the NAcSh network was greater during reward delivery compared to reward omission (Unpaired t-test: Delivery vs. Omission, t _(225597)_ = 161.08, p<0.0001; Figure 5F). This finding indicates that the neuronal activity within the NAcSh network exhibits a more coordinated response during sucrose reinforcement.

##### Reward outcomes reorganized NAcSh functional connectivity patterns

Building on the revealed dense connectivity and small-world architecture of the NAcSh network, we aimed to characterize the strongest functional connections between neuronal ensembles. We quantified the strength of “functional” connections (including self-loops) to identify ensembles with the highest percentage of functional connectivity. While this approach does not provide directionality, it unveils the strongest functional connectivity landscape between neuronal ensembles.

Figure 5G illustrates the functional connectivity pattern for reward delivery. During reward delivery, the 1^st^ strongest functional connections of 13 out of 14 neuronal ensembles were strongly connected with the Choice ensemble, emerging a radial connectivity mode centered on the Ch ensemble (Figure 5G, left panel). Surprisingly, most neuronal ensembles were coordinated by the Ch ensemble, except for the Go ensemble, whose nodes connected more with nodes from the Coh ensemble. The Ch ensemble also exhibited the strongest functional connections with itself, perhaps reflecting a role in facilitating the rat’s integration of information about choice and the outcome obtained, which is needed to update its decision process.

In contrast, during reward omission, the Ch and Om ensembles mainly coordinated the functional connections, followed by the Coh ensemble (Figure 5G, right panel). The Ch ensemble had more functional connections with the Om ensemble, and these two, in turn, connected to the ensemble carrying lick-related activity, the Coh ensemble. Thus, our data suggest that the lick-coherent ensemble is poised to facilitate the integration and synchronization of relevant information in our task.^10^ Collectively, these results unveiled the dynamic reshaping of functional connections between neuronal ensembles in the NAcSh, depending on the reward outcome (see **Figure S10** for complete connectivity matrix).

Having identified neuronal ensembles in the NAcSh and network properties using all 50 trials, we now investigated how their activity changed across the reinforcement learning within a block. We achieved this by repeating the ensemble analysis, focusing on neuronal activity from the Early or Late phases of the task.

#### Neuronal ensembles are dynamic and reorganized by learning

A notable feature of our behavioral task is the requirement that the rat adapt its behavior and reset its reward expectations in each block transition. The rat solely relied on the delivery or omission of the reward as external feedback to guide its choices. Given the importance of this foraging behavior, we aimed to uncover the dynamic changes of neuronal ensembles involved in encoding and processing reward-related information throughout learning.

Compared to the 14 neuronal ensembles identified using all trials, t-SNE, and hierarchical clustering identified 12 and 10 distinct neuronal ensembles using Early and Late trials, respectively (Figure 6A **and Figure S11** for heatmaps). This gradual shrinking in the number of neuronal ensembles explains how each unique activity signature of ensembles consolidates and emerges with learning.

**Figure 6.**
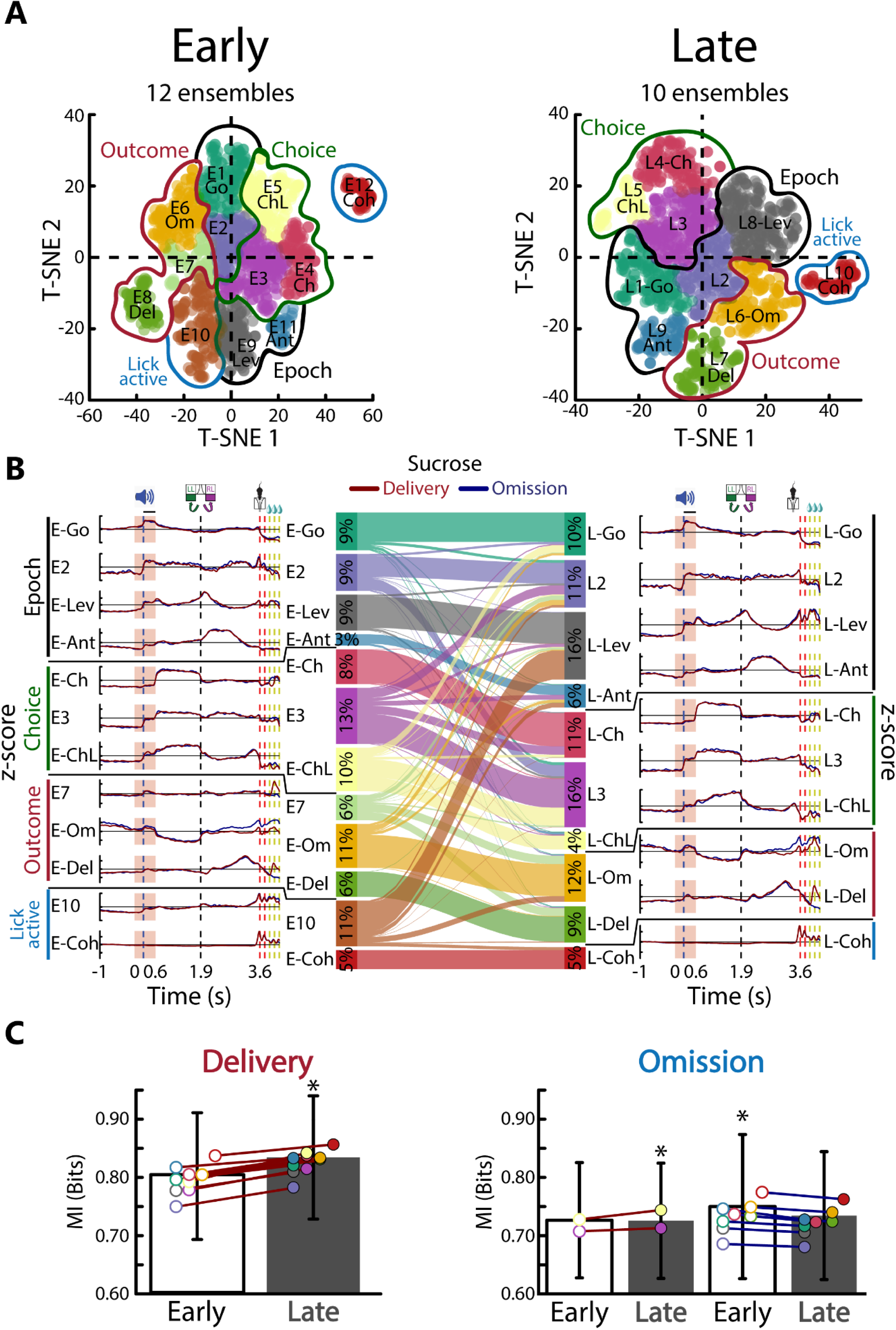
Reinforcement learning consolidates neuronal ensembles and the strength of connectivity (mutual information) during reward delivery and omission. **(A)** t-SNE maps for neural activity in the Early (first 25 trials) and Late (last 25 trials) task phases. The ensembles found in the Early phase were denoted by “E,” while those found in the Late phase were tagged by “L.” **(B)** Left panel, PSTH of average ensemble activity across the 12 neuronal ensembles identified in the Early phase. The red and blue lines represent the mean sucrose delivery and omission activity pattern. In the middle panel, the Sankey diagram draws on how neurons drop in and out between neuronal ensembles from the Early to Late phase (see **Figure S11** for heatmaps) and the percentage of neurons for each ensemble. Right panel, PSTH of the mean activity pattern through the 10 neuronal ensembles found in the Late phase. **(C)** Reward delivery and omission underwent opposite strength connectivity (mutual information) while learning a reinforcement learning task.

Jaccard index was used to track individual neuron transitions between Early and Late phases across neuronal ensembles (**Figure S12**). Thus, we gained insights into the reorganization of these ensembles through learning. Our analysis revealed that nearly all ensembles underwent reshaping, with neurons removed from and added to other ensembles. While most ensembles maintained relative stability through learning (e.g., Coh, Ant, Ch, Del, and Om), some were distributed into other ensembles (E7 distributed into L-Del and L-Om), and others merged (E-Lev and E10 combined into L-Lev). Interestingly, ChL was the only ensemble that reduced the number of neurons through learning (Figure 6B). These findings highlight the stability of some and the dynamic nature of other neuronal ensembles and how their unique activity signature and neuronal composition emerged while solving a reinforcement learning task.

#### Reinforcement learning reorganizes how neuronal ensembles functionally connect

Identifying and characterizing the dynamics of these neuronal ensembles might provide valuable insights into the functional connectivity in the NAcSh network during a reinforcement learning task. We observed that functional connectivity was reorganized by learning during reward delivery and omission (more detail in **Figures S13-14**).

#### Reinforcement learning enhances mutual information during reward delivery and diminishes it by reward omission

Finally, we evaluated how the strength connectivity of the NAcSh network (MI between pairs of neurons) was affected by reinforcement learning. We found that when the reward was delivered, the mutual information significantly increased during the reward-related learning (MI Reward delivery, Unpaired t-test: Early vs Late, t _(225,596)_ = −70.92, p<0.0001; Figure 6C, left panel). In trials where the reward was omitted, the mutual information was not affected uniformly across the neuronal ensembles. Only the ensemble 3 and ChL increased (E3 and ChL MI in Reward omission, Unpaired t-test: Early vs Late, t _(81,133)_ = −9.37, p<0.0001). However, the remaining ensembles decreased significantly (MI Reward omission, Unpaired t-test: Early vs Late, t _(134,102)_ = 28.27, p<0.0001). These results indicate an opposite effect on mutual information in the NAcSh ensemble during reward-related learning and outcome. In sum, the NAcSh network reorganized its functional connectivity pattern, and sucrose enhances network synchrony, whereas reward omission decreases it. The reinforcement learning progressively refined these effects (See Graphical Abstract for a summary).

## Discussion

Our study revealed that neuronal ensembles in the NAcSh act as a specialized module for processing reward-seeking behavior. NAcSh ensembles exhibit remarkable dynamism, progressively reorganizing throughout learning as individual neurons drop in and out, altering the composition of neuronal ensembles, even shrinking in number, most likely reflecting consolidation^34^ and greater similarity in firing patterns. Reward delivery enhances network mutual information, suggesting a coordinated and facilitated flow of reward information. In contrast, omission decreased mutual information. Interestingly, the NAcSh network follows a heavy-tailed distribution, meaning a few highly connected neuronal hubs play a central role in information flow. This architecture most likely facilitates the efficient transmission of reward signals within the network. These findings offer a novel perspective on reward processing in the NAcSh.

### NAcSh Neurons Encode Choice, Outcome, and Licking

Individual NAcSh neurons exhibited diverse activity patterns, representing the rat’s choice (lever selection), reward outcomes (delivery or omission), and rhythmic licking (Figure 2). Some neurons encoded multiple variables, suggesting the NAcSh integrates diverse information beyond tracking reward.^6^

**Choice-selective neurons** could encode choice (or left/right movements), but others could even predict lever choice before the trial began (in baseline). These findings align with the role of the NAc in action monitoring,^35^ action selection,^36^ as well as predicting and guiding actions.

**Outcome-selective neurons,** we observed distinct neuronal populations in the NAcSh selective for reward delivery and omission (**Figure S3A**), suggesting a role in learning from both outcomes. This selectivity was independent of licking behavior, as rats were required to perform a minimum of four licks under both conditions. Under this orofacial controlled condition, we confirmed Regina Carelli’s findings of omission-responsive neurons^7^ and found them equally represented in the NAcSh. PCA analysis further confirmed the presence of distinct outcome-related components (Figure 3), PC3 (reward delivery) and PC4 (reward omission). Our PCA analysis robustly distinguishes orofacial responses from reward signals. Thus, outcome encoding was the second most important variable after licking in the NAcSh. Our findings align with the widespread distribution of reward outcome representations throughout the brain.^10,31,37–41^, The ventral striatum, including the NAcSh, exhibits greater neuronal recruitment than other striatal regions in response to feedback.^36,37^ Given that reward omission can be considered a mild form of aversive stimuli, our results support that NAcSh encodes both appetitive and aversive stimuli.^14,42–44^ Unexpectedly, a novel finding was that while exploring reward signals, we stumble upon neurons that anticipate reward omission, even without explicit cues regarding reward availability (“pessimist” neurons; see also Om ensemble Figure 4D and **Figure S6**). This suggests that the NAcSh may also participate in an “internal” prediction of negative outcomes (reward omission), probably based on reward history (**Figure S5**). A possible implication of having both “pessimist” and reward representations is that the interplay between these two opposing representations poises the NAcSh in a privileged position to compute the motivational balance between cost and benefit to obtain a reward,^44–47^ influencing the willingness of rats to work for sucrose.^48^

**Lick-Coherent neurons and Reward Signals in the NAcSh**, we identified a subset of NAcSh neurons (23.8%) exhibiting strong coherence with licking behavior, consistent with our previous reports.^10,13,15^ These neurons accounted for the most robust responses on NAcSh activity (PC1 and PC2 components). Recent studies using high-density recordings have reported widespread representation of reward outcomes throughout the brain.^31^ They have suggested that such activity may primarily reflect licking movements rather than the hedonic aspects of reward.^31^ While our results confirm the significant role of licking in NAcSh activity, we were able to isolate distinct reward signals from licking itself. Disentangling licking from reward has proven challenging due to their close temporal association. As noted, previous work has shown both anticipatory and suppressed activity in the NAcSh during freely consummatory behavior,^15,18^ supporting that NAcSh inhibition may gate food intake.^3,5,19^ However, separating reward signals from licking under such conditions is difficult due to their inherent overlap. For example, analyzing NAcSh activity under freely licking sucrose intake confounds the approach, the reward, and the oromotor activity, hindering signal separation.^13,15,31,43^ Our novel approach, involving delayed sucrose delivery and enforced dry licks during reward omission trials, allowed us to effectively isolate reward signals. These signals manifested as brief, phasic peaks within one or two licks (see Del Ensemble, Figure 4D), distinct from orofacial evoked responses. This demonstrates the feasibility of parsing reward and licking signals within the NAcSh.

### NAcSh as a hedonic feeding sentinel

Population neuronal activity revealed dynamic modulation of the NAcSh across task epochs (Figure 4E), initiating with the Go signal, followed by an anticipatory peak before licking and a licking-induced inhibition. These findings support the proposed “sensory sentinel” function of NAcSh.^3^ Importantly, we observed that this global NAcSh inhibition, while still above baseline, represents a significant reduction from the robust activation seen during the anticipatory epoch. These results align with the hallmark feature of NAc activity: suppression during palatable food or sucrose intake.^12,15,18,43,49^ Interestingly, we found that the anticipatory peak and the consummatory inhibition in the NAcSh were observed even when the initial licks were empty or during omission trials, in which all licks were dry. This demonstrates that NAcSh inhibition is primarily driven by licking behavior rather than sucrose or palatable food *per se*.^43^

### Neuronal ensembles in the NAcSh are dynamic and serve as distinct modules for reward-related information

Pennartz et al. (1994)^50^ first hypothesized that NAc functions rely on neuronal ensembles of active and inactive neurons. O’Donnell (1999)^42^ suggested that adaptable ensembles, shaped by rewarding glutamatergic limbic inputs,^4,5^ are key to cognition, with reward serving as a driving force for ensemble formation. Building on this seminal work, we present compelling evidence for the dynamic nature of NAcSh ensembles. Extending these classic ideas, we reveal the dynamism of NAcSh ensembles, including their network properties and how their functional connectivity pattern is reorganized with reward outcomes. This underscores the role of dynamic ensembles in understanding the NAcSh function.^9,11^

Our recordings identified several neuronal ensembles encoding reward-related information. We propose that the epoch meta-ensemble encodes specific task epochs that might reveal the current state or the entire trial structure. The Coh ensemble would also be important in defining the outcome epoch (more below). The Ch ensemble was highly informative and not only predicted lever choice (left or right) but also exhibited the greatest percentage of connections of the network (**Figure S8**), suggesting its central role in coordinating functional connectivity during reward delivery (Figure 5G). The high connectivity suggests a potential computational aspect in the Choice ensemble resembling the function of the ventral striatum’s action-value neurons.^36^ We also identified several outcome-related ensembles, such as the Del ensemble, selective for reward delivery. In contrast, Om, E6, E4, Ch, and E2 ensembles showed more robust activation for reward omission. This cornucopia of neuronal ensembles revealed the significant heterogeneity of NAcSh responses and supported the idea that neuronal ensembles are distinct modules specialized in processing specific reward-related information.^22^

### On the complexity of the task and the number of neuronal ensembles

Our study’s use of a cognitively demanding self-guided probabilistic choice task^51^ revealed a striking finding: the engagement of up to 14 distinct neuronal ensembles within the NAcSh, far exceeding the number observed in simpler tasks like freely licking^15^ or brief-access taste tasks.^9^ This suggests a potential link between task complexity and ensemble recruitment, where increased information processing demands may necessitate the activation of additional specialized modules. Alternatively, these complex tasks might unveil a pre-existing, larger repertoire of ensembles that remain dormant during simpler behaviors. Regardless of the underlying mechanism, our findings support the hypothesis that NAcSh neuronal ensembles function as adaptable, task-specific modules. These ensembles dynamically adjust their number and composition based on the specific behavioral context and information processing requirements of reward-guided behaviors.^42^

### NAcSh covariances follow a small-world network architecture with a heavy-tailed degree distribution

Our findings align with Watts and Strogatz’s original “small-world” proposal for the brain’s architecture.^32^ This likely arises from a combination of random connection pruning, synaptic strength rearrangement, and preferential attachment dynamics.^33^ Such mechanisms resemble Hebbian self-organization, suggesting that heavy-tailed connectivity might be a general feature of neural networks.^52,53^ Recent network analyses of the synaptic brain connectome across various organisms (from fly to mouse) support this view, revealing heavy-tailed distributions. Calcium activity within the visual cortex neurons of a single mouse seeing natural images also exhibits this heavy-tailed, “free-scale” property,^53^ echoing our observations in the NAcSh activity. Strikingly, our findings on small-world architecture with heavy-tailed properties in the NAcSh of the ventral striatum align with those observed in brain slices from the dorsal striatum.^54^ Thus, the dorsal and ventral striatum exhibited an asymmetrical tail in the degree distribution between co-active striatal neurons, consistent with a heavy-tailed distribution.^60^

### Licking orchestrates network synchronization

Rhythmic licking unexpectedly emerged as a critical orchestrator of network synchronicity. Across various analyses (PCA, ensemble, network), the Lick-coherent ensemble (Coh) consistently demonstrated a central role. It also exhibits the highest number of neuronal hubs (**Figure S9**). Regardless of reward outcome, the Coh ensemble’s strong functional connections and high mutual information (Figure 5F) suggest a critical function in coordinating neuronal activity, potentially acting as an internal clock synchronizing brain-wide activity.^10,39,55^

### Implications for neuroscience: Can we catch functional ensembles?

The next challenge in neuroscience is to map how genetically defined cell types correspond to “functional” neuronal ensembles based on neural activity. The Allen Brain Institute recently unveiled 5322 distinct cell types in the mouse brain,^56^ raising the question of how these cell types relate to the “functional” neuronal ensembles. Perhaps each functional ensemble consists of only one cell type, or ensembles are composed of multiple cell types sharing a similar firing pattern.^57^ Evidence supports both possibilities,^9,11,14,15,57^ but, given that a single cell type may play multiple functions, it is more likely that various cell types composed a “functional” ensemble.^58^

New techniques like Fos, Arc TRAP^59^, Cal-Light^60^, or mArc^61^ offer promise in identifying activated neurons, potentially capturing “functional” ensembles or even an engram that can induce a percept or memory.^62,63^ However, challenges remain, e.g., gradual increases in activity throughout the trial, from Go cue to reward outcome, making isolating specific ensembles difficult.^39,41^ Suppose we want to capture neurons based on activity and time window (e.g., within the Choice epoch). In that case, capturing selectively, the Ch ensemble will also be trapped with other ensembles that fire more during the Choice epoch (i.e., A3, Lev, A4, ChL, A6, A9) and with other active neurons depending on the trial type (e.g., neurons responding to left-right choice, rewarded or not). Moreover, given that neuronal ensembles are dynamic,^64^ and thus, their neuronal composition changes with learning, as we and many others suggest.^34^ In that case, the neurons trapped will depend on the learning progress of the task. An even greater challenge will be distinguishing neuronal hubs from other non-hub cells within the same functional ensemble due to their similar response patterns (**Figure S9D**). Our results predict that these impressive new techniques may still lack the necessary precision to dissect and fully understand the complexity of functional neuronal ensembles.

### What is the cellular identity of neuronal hubs?

Our network analysis revealed a “rich get richer” connectivity pattern, resembling networks with preferential attachment.^33^ This architecture facilitates rapid information transfer and is resilient to random deletions (removal of weakly connected neurons). However, it becomes vulnerable when explicitly targeting the highly connected “hubs.”^65^ This raises intriguing questions: who are these hubs? (see **Figure S9**) Are they specific cell types, like interneurons?^21,65^ If so, they constitute less than 10% of NAcSh neurons.^66^ Studies in brain slides from the dorsal striatum suggest that most hub neurons are, in fact, interneurons, fast-spiking, low threshold spiking, and cholinergic interneurons.^65^ Alternatively, could they be medium spiny neurons (MSNs, comprising ∼90% of NAcSh neurons)? If so, what distinguishes hub MSNs from other non-hub MSN neurons? Future studies should explore these fascinating avenues to illuminate these critical network elements’ nature and functional significance.

### Limitations of the study

Here, we used PCA and t-SNE followed by hierarchical clustering, to define “functional” neuronal ensembles. While different dimensionality reduction and clustering methods like UMAP,^67^ CEBRA,^68^ or graph theory^65^ might yield slightly different ensemble numbers, the fundamental observation remains consistent: neuronal ensembles emerge^9,11^ as important players in brain computations, regardless of the specific mathematical approach.^22^ Likewise, our correlation matrix leverages data from multiple sessions and rats, capturing the strength of “functional” connectivity related to task events in a kind of “meta-brain” of 17 animals performing the same behavioral task, and thus, it lacks individual-level detail because of restrictions imposed by current recording methods.^31,38,41^

## Acknowledgments

Benjamin Arroyo is a doctoral student from the Programa de Doctorado en Ciencias Biomédicas, Universidad Nacional Autónoma de México (UNAM), who received a fellowship from CONAHCyT, México. The data in this work is part of his doctoral dissertation in the Posgrado en Ciencias Biomédicas of the Universidad Nacional Autónoma de México. We thank Mario Gil Moreno for building homemade electrodes. We also thank Fabiola Hernandez Olvera for her invaluable animal care.

## Author Contributions

BA and RG designed the research. BA performed the research. BA, EHL, and RG analyzed the data, BA, EHL, and RG wrote the paper. All authors reviewed and approved the manuscript for publication.

## Supplemental information

Document S1

Figures S1-S14

Video S1, PCA trajectories as a function of neuronal ensembles, related to Figure 3

## Ethics Statement

CINVESTAV Animal Care and Use Committee reviewed and approved the animal study.

## Methods

### Experimental model and study participant details

Seventeen male Sprague-Dawley rats (300-350 g) were individually housed in a temperature-controlled room (22 ± 1 °C) with a 12-hour light/dark cycle (lights on at 06:00, off at 18:00). The rats were water-deprived and only had access to water for 30 minutes after each behavioral session. Chow food (PicoLab Rodent Diet 20, MO, USA) was always available *ad libitum*. The Animal Care and Use Committee of CINVESTAV approved all procedures.

### Apparatus

Training and recording were conducted in a standard operant conditioning box (30.5 x 24.1 x 21.0 cm; Med Associates Inc., VT, USA). The box had a front panel with a central V-shaped port equipped with an infrared sensor to detect head entry. Two retractable levers were also placed on the sides of the central port. On the opposite wall was a V-shaped licking port with an infrared sensor to detect individual licks. This port contained a licking spout connected to a solenoid valve (Parker, OH, USA). The spout delivered a sucrose solution (10% w/v sucrose dissolved in distilled water). The volume of each drop was calibrated daily to ∼10 µL using an independent air pressure system.^10^

### Behavioral task

A self-guided probabilistic choice task was used,^23,25^ also known as a 2-armed bandit. The training period for the trial-and-error task lasted from 30 to 60 days and included a first pretraining phase (2 days to 2 weeks) before the rats progressed to the entire task.

### Pre-training phase

To complete the pretraining phase, rats had to complete at least 150 trials. Each trial began with the house lights illuminated, which instructed the rat to approach the central port and place its head for at least 100 ms. Once this fixed interval elapsed, an auditory stimulus was delivered, and a single lever was extended. The lever alternated between the left and right lever on consecutive trials (e.g., if the left lever was extended in the current trial, the right lever would be extended in the subsequent trial). After the lever was pressed, the rat had to approach the reward port on the opposite side of the box. The rat was required to deliver two dry licks to the spout, and three drops of sucrose were delivered in the subsequent three licks. All trials were rewarded in this pretraining phase, and sessions were limited to 30 minutes.

### Complete task

In the complete task version, the rat was required to wait 250 ms before an auditory stimulus was presented. Both levers were then extended into the box. The auditory stimulus prompted the rat to perform a leftward or rightward lever press but did not provide guidance on which lever would deliver the highest reward. Reward probabilities were assigned to the left and right levers, covering all combinations of 10%, 50%, and 90%. The reward blocks changed every 50 trials without an explicit signal, and the order of blocks varied across sessions.

After the lever press, both levers were retracted simultaneously to prevent the possibility of a second press in a single trial. The rat then had to approach the reward port on the opposite side of the box and give two dry licks to the spout, regardless of the outcome. In rewarded trials (reward delivery), the spout delivered a drop of sucrose in the following three licks. In unrewarded trials (reward omission), the rat had to give two more dry licks to finish the trial (i.e., at least 4 dry licks). Trials ended 0.5s after the last lick was given in the reward port. No external cues could influence the rat’s choices aside from the outcomes of its actions.

A single trial structure of self-guided probabilistic choice task: A trial began with the lights on, and the box illumination instructed rats to nose poke the central port. An auditory tone (880 Hz and 80 dB) was delivered if the rat remained in that position for 0.25 s. The Baseline epoch comprised −1 to 0.25 s from head entry. At the onset of the Go signal, two levers were extended, and the rat headed out from the central port (Go epoch). In the Choice epoch, rats were required to choose and press one of the two levers (left-green or right-purple color). Both levers were retracted after each choice, and the Anticipation epoch began. In this epoch, rats needed to approach the reward port on the opposite side of the box. During the Outcome epoch, when a trial was rewarded, after two dry licks, a drop of 10% sucrose was delivered in the following three licks, one drop per lick (Delivery, see the three red L’s and the three drops). Otherwise, the reward was omitted (Omission trial). The trial was finished 500 ms after rats refrained from licking and the light was turned off. From block to block, rats had to infer the reward probability associated with each lever to self-guided choices and update their beliefs every 50 trials. The transition between blocks was not cued.

### Surgery

Animals were anesthetized with ketamine (70 mg/kg, i.p.) and xylazine (20 mg/kg, i.p.). They were then placed in a stereotaxic apparatus, where the skull was exposed, and craniotomies were made in the skull over the recording site. Two screws were set for holding, and a silver wire was soldered to the third screw. This served as an electrical reference for recordings and was inserted above the cerebellum. A custom-made 16-tungsten electrode array (35 µm diameter, each arranged in a 4 x 4, area = 1 mm²) was implanted into the NAcSh (AP: +1.2 mm, ML: ±1 mm from bregma; DV: −7.6 mm). The electrode array was stained with a fluorescent dye (cell tracker CM-Dil dye, red excitation/emission spectra: 553/570 nm; ThermoFisher Scientific, MA, USA) just before it was inserted into the brain. Dental acrylic was applied to the screws, making a stable attachment to the skull and fastening the electrode array. The rats were given ketoprofen (45 mg/kg, i.p.) three days after the surgery and recovered for one week.

### Histology

Once electrophysiological signals disappeared, the animals were deeply anesthetized with an overdose of pentobarbital sodium (150 mg/kg, i.p.). They were then transcardially perfused with phosphate-buffered saline (PBS 1x) followed by 4% paraformaldehyde. Brains were extracted, stored for 1 day in 4% paraformaldehyde, and then transferred to a 30% sucrose/PBS solution. Brains were cut at 60 µm coronal slices. An epifluorescence microscope confirmed the trace and last position of the electrode array tips.

### Electrophysiological recordings

Recordings were made while rats performed the probabilistic choice task. Neural activity was amplified, digitized with a 64-channel PZ5 Neurodigitizer amplifier (Tucker-Davis Technologies, FL, USA), and then transmitted to the RZ2 BioAmp Processor (Tucker-Davis Technologies, FL, USA). Signals were sampled at 50 KHz (28 bits of resolution) and band-pass filtered from 0.5 to 8 KHz. The action potentials were identified online using voltage-time threshold windows. To be classified as a single-unit, action potential needed to have an amplitude above 50 µV, and their spikes had to exhibit an amplitude of 3.5-4 standard deviations above the mean signal of the channel. The spikes were exported (Open Bridge software, Tucker-Davis Technologies, FL, USA) and sorted using Offline Sorter Software (Plexon, TX, USA). Only time stamps from the offline-sorted waveforms were analyzed.

### Reinforcement learning model

A Q-learning model was employed to predict the decision-making behavior (probability of choosing the left lever, p(Left-Lever)) by leveraging subject-specific reward history and action sequences to estimate action value. Action-values were calculated on a trial-by-trial basis and updated to the following algorithm:

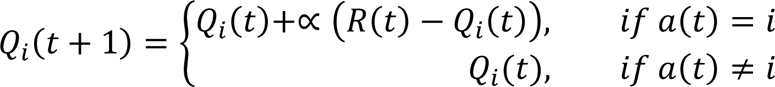

Where *a*(*t*) and *R*(*t*) indicate the choice and reward in the current trial, respectively, *i* is the action associated with the value (e.g., left or right lever), and ∝ is the learning rate. This model states that in two alternative task, the probability of choosing an action is a sigmoidal function of the difference in the action values:

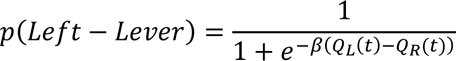

Where β is a parameter that determines bias towards the action associated with the higher action value. The learning rate and bias parameters were estimated from the behavior for each session as in reference ^69^.

### Time-warping algorithm

We determined 5 epochs in the task: 1 s before the head entry to the onset of the tone (Baseline), from the onset of the tone to head exit (Go), from the head exit to the press lever (Choice), from the press lever to the first lick (Anticipatory), and from the first to the fifth lick (Outcome). These epochs were aligned by implementing the time-warping algorithm, given the variability of the time intervals across trials. This algorithm was not applied to the Baseline epoch. The epochs were aligned by calculating the median of the time intervals (**Figure S2**) for Go = 0.35 s, Choice = 1.3 s, and Anticipatory = 1.9. The outcome epoch was aligned lick-by-lick. Thus, we calculated the Inter-Lick Interval (ILI) median = 0.14 s. The duration of an epoch was calculated in each trial:

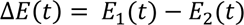

where *E*_1,2_ (*t*) are the events which start and terminate the epoch, respectively, and *t* is the trial. We defined an upper limit (*UL_Epoch_)* for each epoch duration given its cumulative density function. If Δ*E*(*t*) exceeds its corresponding epoch limit, Δ*E*(*t*) was determined following this formula:

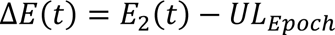

We considered these trials as delayed, avoiding the over-shrink epoch. To set the spike timing in the same interval of the duration of the epoch was given by:

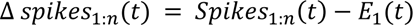

Or when the epoch duration exceeds its upper limit:

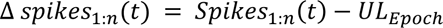

Where *Spikes_1:n_(t)* are the spike timestamps in the epoch. A range normalization of the epoch was calculated as follows:

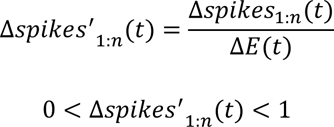

Finally, the spike timing in the epoch of each trial was linearly transformed into its corresponding median of the epoch:

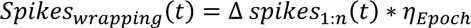

Note that the algorithm enables us to preserve the number of spikes and the proportion of the inter-spike interval. The Time-warping algorithm shrinks or expands the temporal scaling by conserving the pattern and amplitude of the firing rate of the neurons.

In addition, we considered evaluating the neural activity until the fifth lick for the outcome epoch, as this parameter was found to be consistent across most trials. Including this task condition was considered important to perform a fair comparison for the subsequent neural analysis, particularly for those in which reward was delivered versus those in which it was omitted. By ensuring that both types of trials were subject to the same task constraints, we conducted a more rigorous comparison of the licking behavior on both reward outcomes.

### The neuronal responses of the NAcSh

Following the time-warping algorithm, the firing rate of the 1507 NAcSh neurons recorded was calculated using the convolution with a Gaussian kernel (σ = 30 ms):

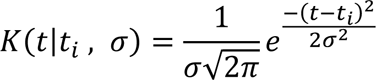

Where (*t*−*t_i_*) represents a spike occurring at the time *t_i_* (i.e., timestamps of every spike per neuron). The estimate of the firing rate given by the convolution was:

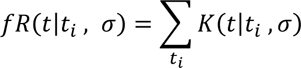

Then, we normalized neuronal responses using a z-score:

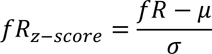

Where *fR* was a time series of the firing rate of a given neuron along the trial, corresponding to 1 s before the onset of the head-in and the fifth lick delivered onto the spout in the reward port (4.16 s after the onset of the head-in). The mean and standard deviation were calculated from the firing rate within the trials.

### Outcome and Choice identification NAcSh neuronal responses

NAcSh neurons have diverse pattern responses associated with different task variables, such as Choice (Left or Right lever) or Outcome (Delivery or Omission). The Receiver Operating Curve (ROC) analysis was implemented to quantify how different the neural firing rate for Choice (i.e., firing rate distribution during left-lever trials vs. firing rate distribution during right-lever trials) and Outcome (i.e., firing rate distribution during reward Delivery trials vs. firing rate during reward Omission trials) over time (i.e., number of consecutive bins, the bin width was systematically adjusted from 100 to 1000 ms in increments of 100 ms. Then, we chose the bin width, which reduced close to zero the proportion of neurons that were selective to both conditions (e.g., reward Delivery and Omission trials) in different time intervals. Thus, given these criteria, the optimal bin-width was 400 and 200 ms for choice and outcome, respectively.

### Lick coherent neurons

Neuronal activity of the NAcSh exhibited spiking responses consistent with rhythmic licking behavior (within the bandwidth of 4 to 12 Hz). This coherence was identified using a multi-taper time-frequency coherence analysis.^70^ The confidence intervals and the significant threshold were determined using a jack-knife method. Neuronal responses were considered lick-coherent if the lower confidence interval was above the significant threshold for only delivery, omission trials, or both (delivery & omission).^10^

### Principal Component Analysis

The Principal Component Analysis (PCA) was implemented to understand the neural dynamics of the NAcSh population responses. Following the time-warping algorithm and the preprocessing, the neuronal responses were concatenated in an n x m matrix. Each row was the mean firing rate of a single unit, and columns were the time bins at 5 ms. The mean neuronal responses were vertically concatenated for 3 types of trials: all trials (Average), only trials where the reward was delivered (Delivery trials), and only trials where the reward was omitted (Omission trials). Linear combinations of population activity were computed, capturing the most variance in the averaged population responses. The population activity was projected onto three axes, describing the population trajectories across the task’s epochs.

### Ensemble classification

Using the same n x m, a concatenated matrix was used to classify them into clusters or, hereafter referred to as ensembles. Considering the high dimensionality of the concatenated NAcSh neuronal responses, the PCA was implemented prior to the t-distributed Stochastic Neighbor Embedding (t-SNE) using the first ten principal components embedding points taking into consideration to preserve neighborhood identity optimally.^71^ Using the “Elbow curve” method, the number of clusters for the three-block phases (i.e., All, Early, Late) to optimize was set to 20, and the number of ensembles found for each block condition was 14 for All, 12 for the Early phase, and 10 for the Late phase. Using the number of ensembles optimized for each block phase, neurons were grouped using a hierarchical clustering analysis, which uses the increase in the total within-cluster sum of squares of the distances between all points in the cluster and the centroid of the cluster. The neuronal responses were grouped using a similar method as in ^9^.

### Methods Network inference and analysis

Probabilistic graphical models (Markov random fields) were inferred from the electrophysiological recordings by calculating the mutual information (MI), an information theoretical measure of statistical dependency (MI is the maximum entropy or less-biased estimate of statistical dependence) using the infotheo: ‘information-theoretic measures: R package version 1.2.0’.^72^ After binning discretization, MI (in bits) was calculated between all neurons recorded per experiment. Once the full N x N adjacency matrix was obtained (with N the number of recorded neurons per experiment), an approach to graph sparsification was achieved through quantile thresholding. The 10% higher value of the distribution tail of the mutual information was determined, and all links equal to or above this threshold were retained while discarded elsewhere. The set of links and nodes retained form a probabilistic network for each condition.

Networks were visualized, and their resulting topological structure was analyzed using Cytoscape^73^ to determine local and global centrality measures. Connectivity degree distribution, local and global clustering coefficients, average shortest path length, and betweenness centrality distributions were also determined using the Network Analyzer package in Cytoscape.

### Quantification and statistical analysis

All analyses were performed with Matlab (Mathworks) and R (http://www.r-project.org/). An unpaired t-test was employed to evaluate the impact of surgery on choice behavior, biases in lever selection based on reward probability, and differences in lick distribution during reward delivery and omission. Kruskal-Wallis test compared the stay or shift probability as a function of the number of rewards delivered or omitted in the last seven trials and the probability of pressing the left lever and reward obtained across blocks. We calculated the chance threshold to assess the significance of the fraction of neurons identified using the ROC method for detecting Outcome or Choice-selective neurons. This threshold represents the number of neurons considered significant (using the same ROC criteria) purely by chance. To establish this threshold, we performed shuffling experiments on the outcome and choice variables (delivery vs. omission, left vs. right, respectively). In each iteration, we shuffled the trials, calculated the ROC, and repeated this process 1,000 times. The chance threshold for each variable was then determined based on the fraction of neurons that achieved significance through random chance. Chi-square tests were employed to assess significant hemispheric lateralization of right-selective neurons. This test was also employed to assess significant differences in the fraction of neurons selective to specific variables (e.g., Outcome, Choice, Coherence) or between fractions of selective neurons (e.g., Delivery-selective vs. Omission-selective) across neuronal ensembles. Jaccard index was used to quantify the overlapping proportion of neurons transitioning between block conditions (i.e., All, Early, Late) across neuronal ensembles (using jaccard.test() in “R”). Unpaired t-test compared the difference in mutual information of functional connections between neuronal ensembles across reward outcomes (e.g., reward delivery or omission) and block conditions (e.g., Early, Late). All data are presented as mean ± SEM unless reported otherwise. The accepted level of significance was a p≤0.05.

