## Supplemental Figures for "The flow of reward information through neuronal ensembles in the accumbens"

**Supplementary Figures**

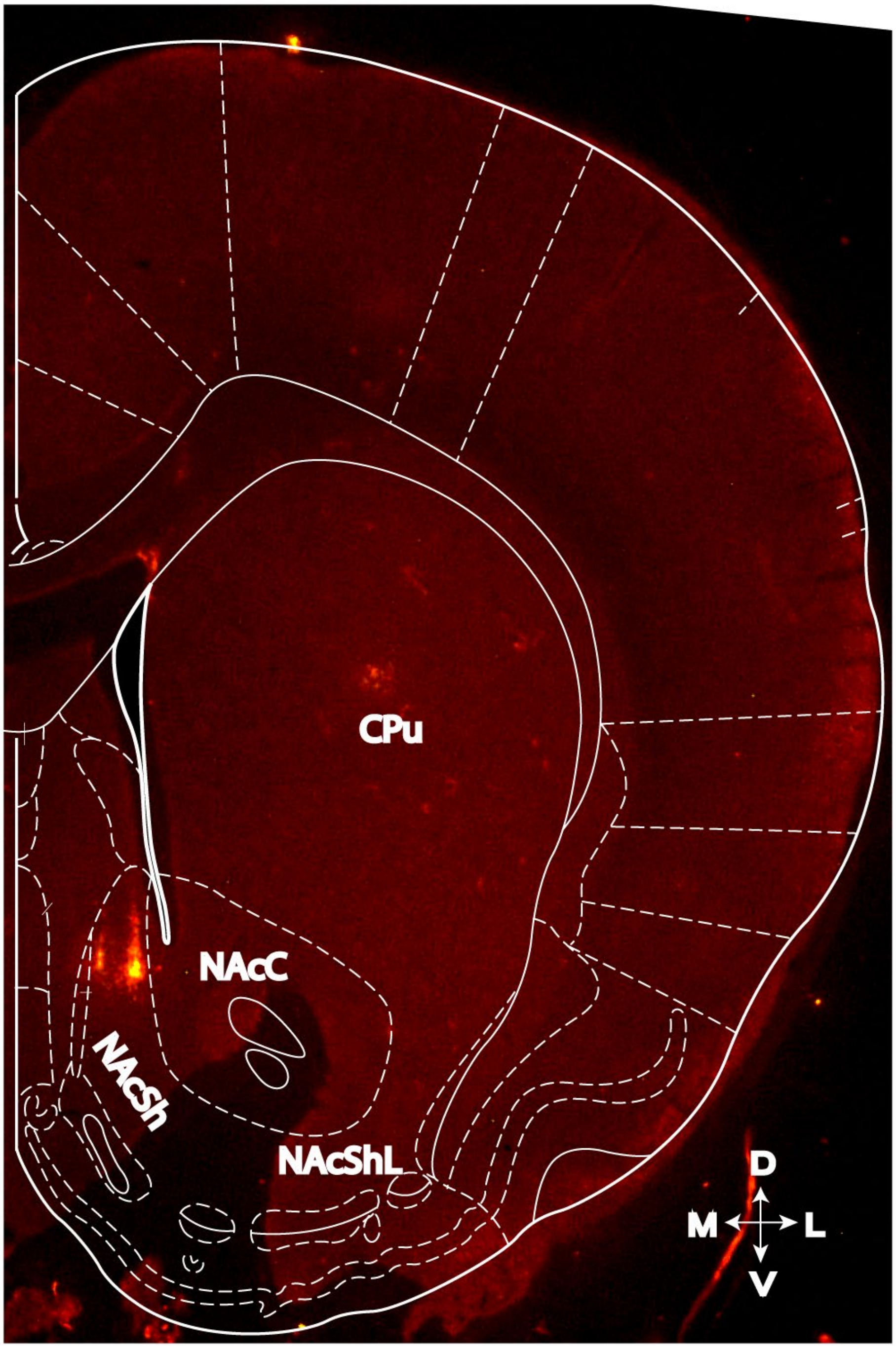


**Figure S1. A representative track of recording sites.**

A coronal mouse brain section was viewed in an epifluorescence microscope in which the multielectrode array was stained by a fluorescent dye (cell tracker CM-DiI dye, red excitation/emission spectra: 553/570 nm; ThermoFisher Scientific, MA, USA). The electrode array was found in the anterior and medio-dorsal parts of the Nucleus Accumbens Shell (NAcSh). NAcC = nucleus accumbens core and NAcShL = nucleus accumbens lateral. CPu, = Caudate putamen

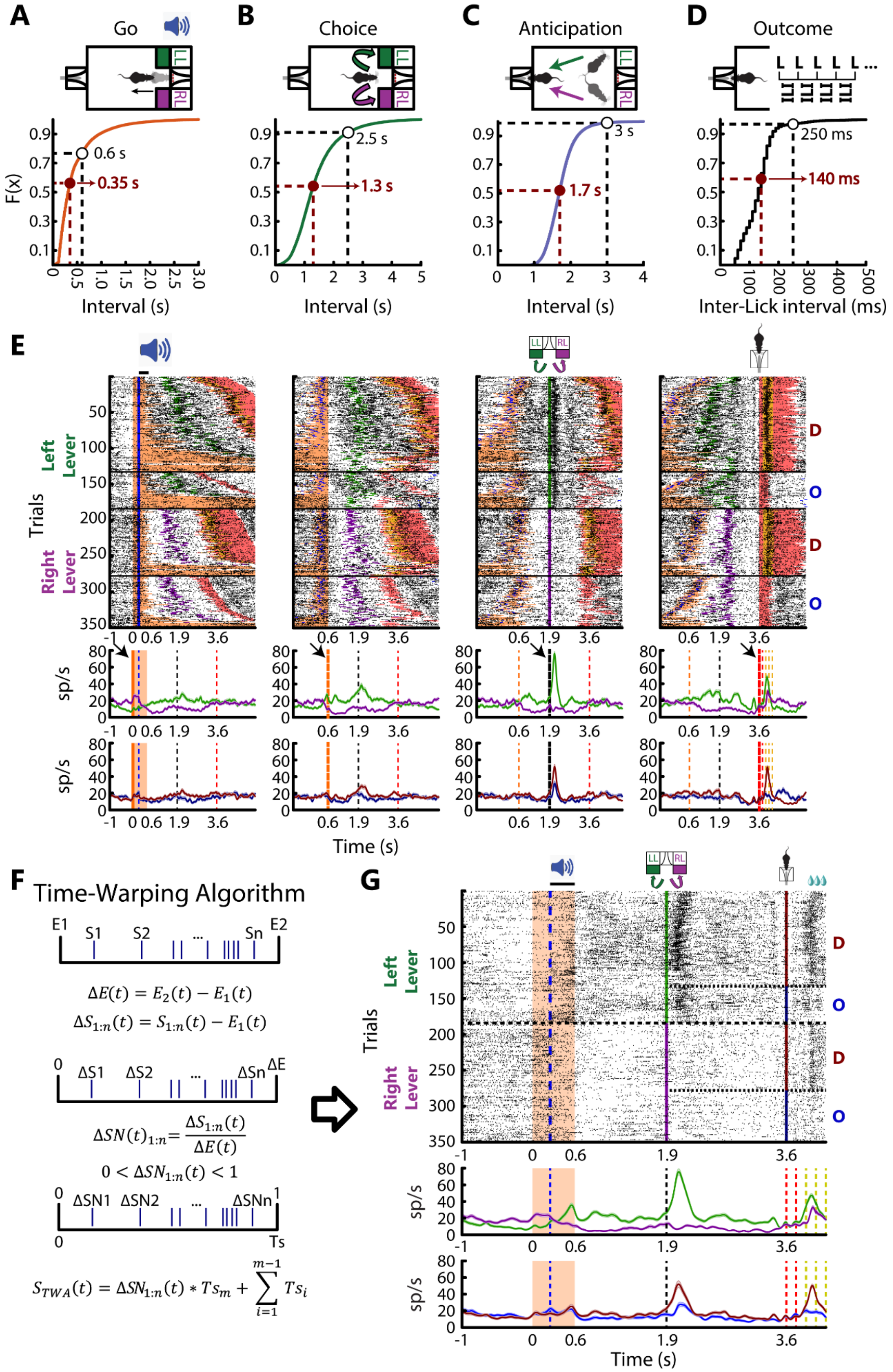
A
Go
B
Choice
C
Anticipation
D
Outcome
F(x)
0.6 s
0.35 s
2.5 s
1.3 s
3 s
1.7 s
250 ms
140 ms
Interval (s)
Inter-Lick interval (ms)
E
Trials
Left Lever
Right Lever
D
O
sp/s
Time (s)
F
Time-Warping Algorithm
E1
S1
S2
Sn
E2
$\Delta E(t) = E_2(t) - E_1(t)$
$\Delta S_{1:n}(t) = S_{1:n}(t) - E_1(t)$
ΔS1
ΔS2
ΔSn
ΔE
$\Delta SN(t)_{1:n} = \frac{\Delta S_{1:n}(t)}{\Delta E(t)}$
$0 < \Delta SN_{1:n}(t) < 1$
ΔSN1
ΔSN2
ΔSNn
Ts
$S_{TWA}(t) = \Delta SN_{1:n}(t) * Ts_m + \sum_{i=1}^{m-1} Ts_i$
G
Trials
Left Lever
Right Lever
D
O
sp/s
Time (s)

**Figure S2. Representative raster-plot of a time-warped neuronal response.**

**(A)** Top panel illustrates the Go epoch, the reaction time between the Go signal and exit of the head from the central port. Below, the cumulative distribution function of these time intervals is presented. Dashed red lines show the time constant used for alignment (close to the median of the distribution), with dashed black lines indicating the upper limit (about 1.8 times the median) to prevent an over-shrink effect (see methods).

**(B-C)** Same convention as A, but for Choice and Anticipatory epochs.

**(D)** Depicts the distribution of inter-licks intervals (ILI) observed in the Outcome epoch.

**(E)** A raster plot of an example neuron pre-time-warping. Spiking timestamps (black ticks) of a NAcSh neuron are aligned (time = 0s) to four task epochs: head entry, head out, lever press, and the first empty lick on the outcome spout. The orange-shaded area represents the head-entry time interval in the central port. The vertical blue line indicates the Go signal onset, and the green and purple lines show the left and right lever press times, respectively. The shaded red area denotes the time of licking duration in the reward port. The yellow mark indicates the three drops of sucrose. Below is the PSTH of trials sorted by left/right choices (top PSTHs, green and purple lines, respectively) and reward delivery/omission (bottom PSTHs, red and blue lines, respectively). The orange area depicts the time interval in the central port. The black dashed line indicates the onset of the lever press, the dashed red and yellow lines indicate the times at the first two empty licks. The three licks associated with sucrose were delivered or omitted in the reward port, respectively. The arrows indicate the time for alignment.

**(F)** A graphical depiction of the Time-Warping Algorithm (TWA).

**(G)** A Raster plot of the same neuron is shown in **E** after TWA. This algorithm linearly scales the time intervals between task events on a trial-by-trial basis approximately to the median interval, preserving the number of spikes and overall firing pattern between events.

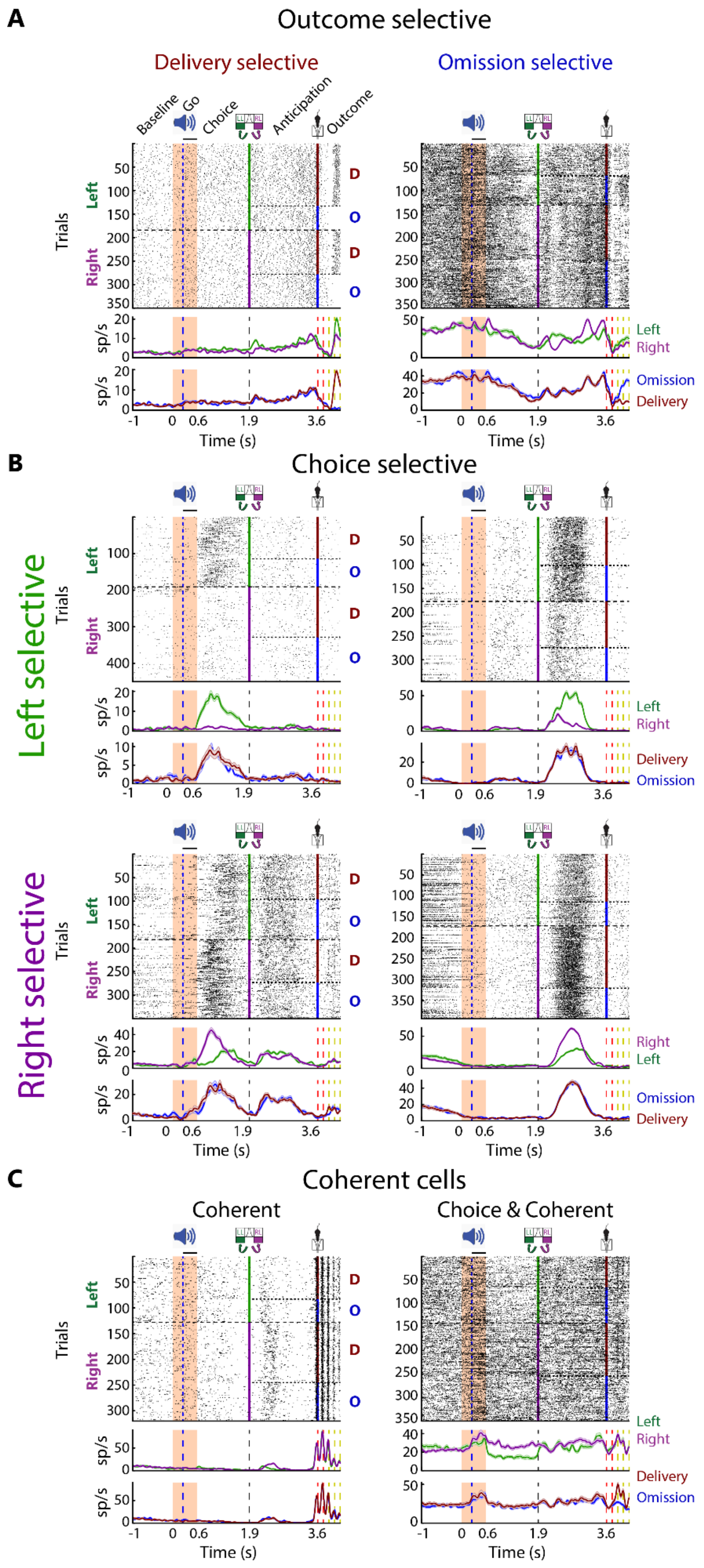
A
Outcome selective
Delivery selective
Omission selective
Baseline
Go
Choice
Anticipation
Outcome
Trials
Left
Right
D
O
sp/s
Left
Right
Omission
Delivery
Time (s)
B
Choice selective
Left selective
Right selective
C
Coherent cells
Coherent
Choice & Coherent

**Figure S3. More examples of individual neuronal responses of the NAcSh.**

The neuronal responses are shown in the same convention as in **Figure 2A-C**.

**(A)** NAcSh outcome selective neurons. The left panel depicts a representative delivery-selective neuron that increases its firing rate specifically when a reward (sucrose) is delivered but remains silent when it is absent. In contrast, the right panel shows an omission-selective neuron activated only when the reward is omitted.

**(B)** NAcSh choice-selective responses. The upper panels depict two neurons with an increased firing rate selectively during the activation of the left lever. Conversely, the bottom panel presents two neurons whose firing rates become selectively increased in response to the right lever. The left panel is a neuron whose firing rates are selective to the respective lever during the Choice epoch, whereas the right panels are selective during the Anticipation epoch.

**(C)** A lick-coherent neuron in the NAcSh. In the left panel, a representative neuron whose firing rate is coherent to licking, while in the right panel, a neuron whose firing rate is coherent only during the sucrose delivery but not omission. The lick-coherent neuron at the right panel also exhibited a choice-selective response during the Choice epoch.

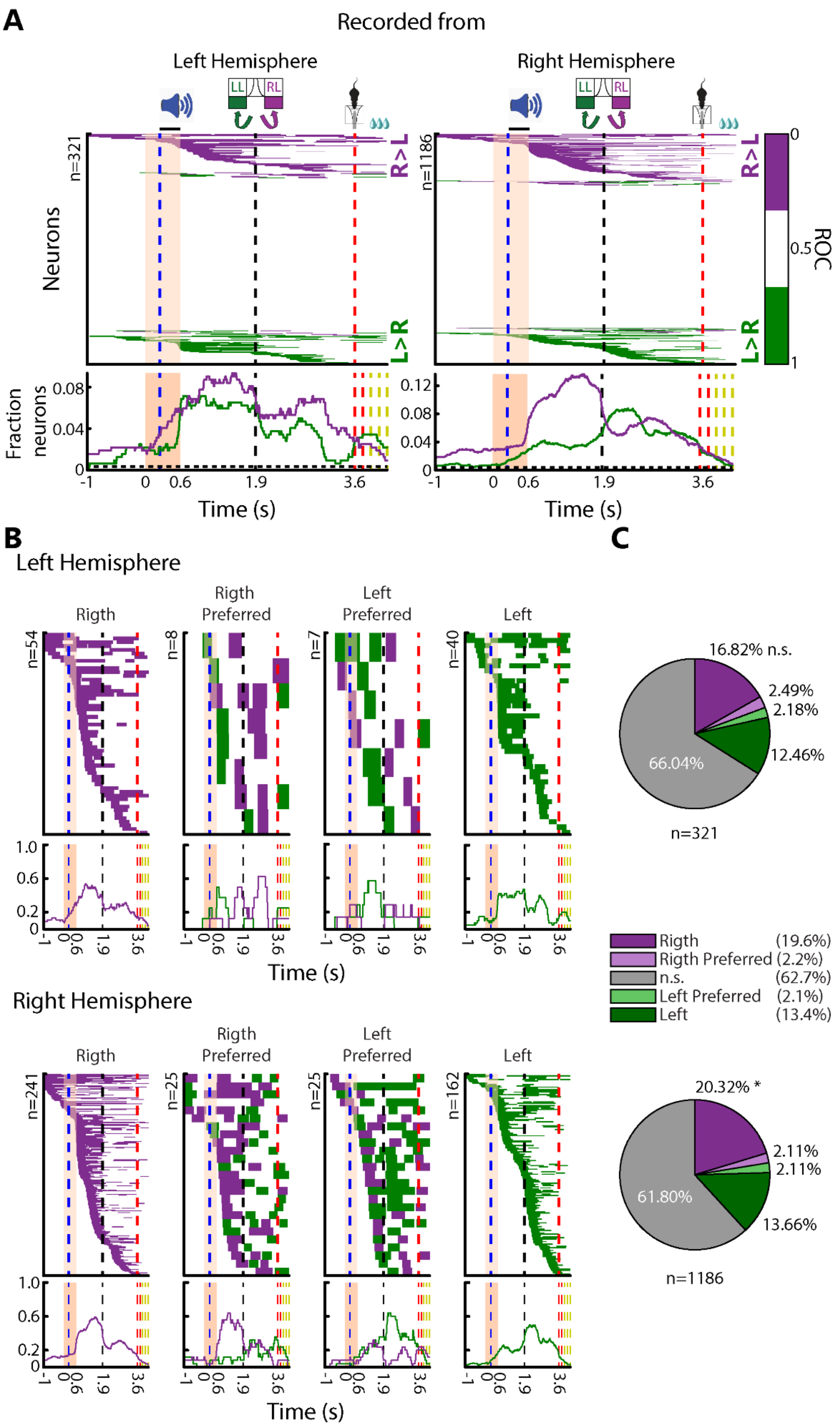
A
Recorded from
Left Hemisphere
Right Hemisphere
LL
RL
n=321
n=1186
Neurons
R>L
L>R
ROC
0
0.5
1
Fraction neurons
Time (s)
B
Left Hemisphere
Rigth
Rigth Preferred
Left Preferred
Left
Right Hemisphere
C
16.82% n.s.
2.49%
2.18%
12.46%
66.04%
n=321
Rigth (19.6%)
Rigth Preferred (2.2%)
n.s. (62.7%)
Left Preferred (2.1%)
Left (13.4%)
20.32% *
2.11%
2.11%
13.66%
61.80%
n=1186

**Figure S4. While right-lever selective neurons were present in both hemispheres, the right hemisphere contained a significantly higher proportion of these neurons.**

**(A)** Neurons were identified by ROC curve consistently activated by lever press throughout the trial, recorded from the left and right hemispheres (left and right panels, respectively). Top panel, neurons are displayed for each row, purple or green colored bins indicate significant activity for the right or left lever press, respectively. Neurons are sorted by the onset latency of the significant activation. Bottom panel, the temporal fraction of neurons was selectively activated by the lever press throughout the trial. The horizontal dashed line indicates the chance threshold (see Methods).

**(B)** Four different groups of neurons were selectively activated by the lever press throughout the trial for neurons recorded from the left and right hemispheres (top and bottom panel, respectively).

**(C)** The percentage of neurons recorded in the left and right hemispheres (top and bottom pie charts, respectively). A significantly greater fraction of right-selective neurons compared to the left-selective neurons was found in the right hemisphere (20.32% vs. 13.66%, $\chi^2=18.18$, $p<0.0001$), but no significant difference was found on neurons recorded from the left hemisphere (16.82% vs. 12.46%, $\chi^2=2.11$, $p=0.1267$).

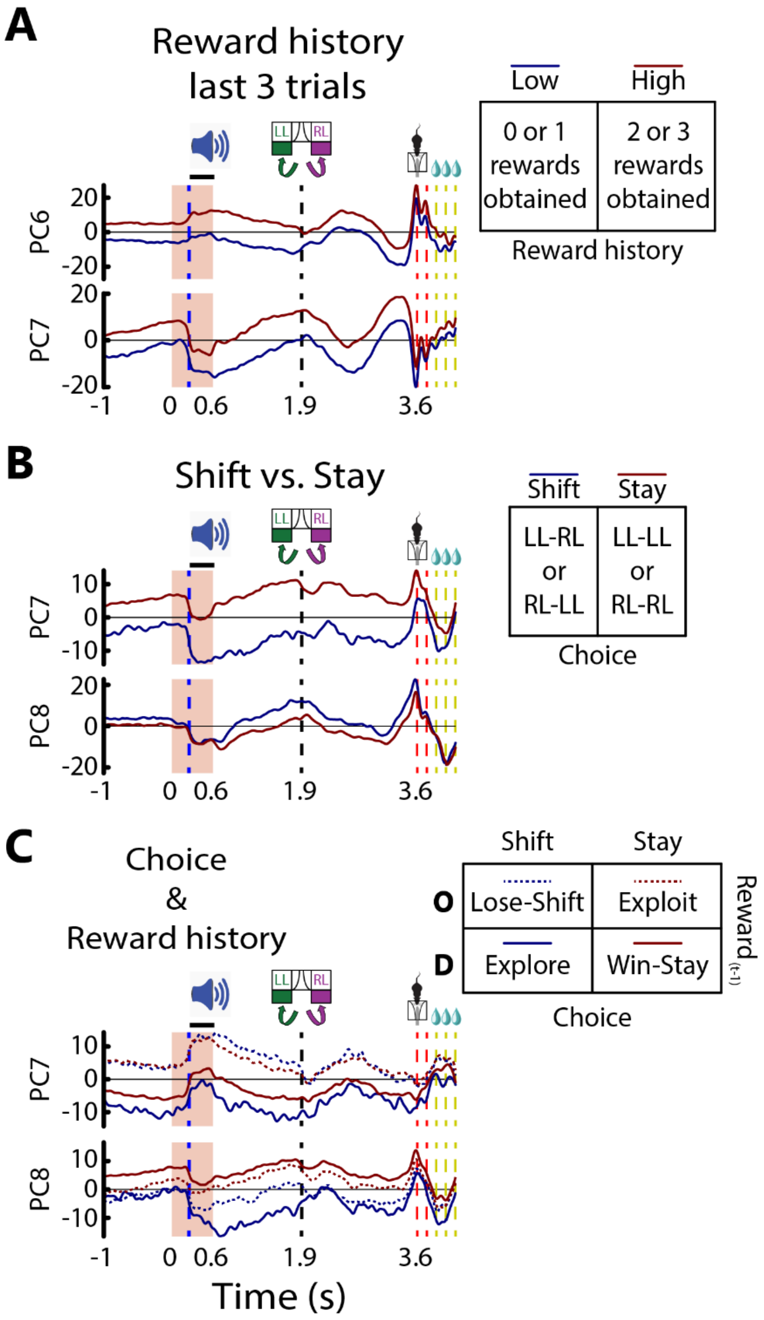


**Figure S5. NAcSh population activity encodes reward history and choice strategy.**

**(A)** PCA analysis reveals distinct population trajectories in the NAcSh (n=1507 neurons) based on reward history (last three trials). The neuronal firing increased when rats received multiple rewards (red) compared to when they received few or no rewards (blue), particularly in PC6 (4.27%) and PC7 (4.15%). These PCs also exhibit activity related to licking behavior. The trajectories of PC6 and PC7 suggest that the NAcSh activity reflects information about the rat's reward history and how successfully it has recently obtained rewards.

**(B)** Further analysis incorporating the rat's choice strategy reveals additional separation in population trajectories. PC7 (3.41%) exhibits higher activity when the rat stays on the same lever (red line), whereas PC8 (2.84%) shows higher activity when the rat switches levers (blue line). The inset illustrates two trial types: Stay trials, when the rat presses the same lever as in the previous trial (e.g., Left Lever-LL, Right Lever-RL), and Shift trials when the rat switches to the opposite lever (e.g., LL-RL, RL-LL).

**(C)** Combining reward history and choice strategy further differentiates population trajectories. PC7 (2.67%) primarily reflects reward history, with higher activity after reward (solid lines) versus no reward (dashed lines). PC8 (2.23%) encodes the rat's choice strategy, with higher activity for staying (red lines). In the inset, "O" depicts omission, "D" delivery in the previous trial (Reward $_{(t-1)}$), whereas Shift indicates if the rat changed lever and Stay if kept pressing the same lever in the current trial. Thus, in this particular analysis, the animal "Explore" when the previous trial was rewarded, but it shifted the lever, and "Exploit" when the reward was omitted in the previous trial but stayed on the lever.

Importantly, these distinct population trajectories encoding reward history and choice strategy are evident throughout the entire trial, from baseline to anticipatory epochs, but rapidly converge during the outcome epoch when the reward is delivered or omitted.

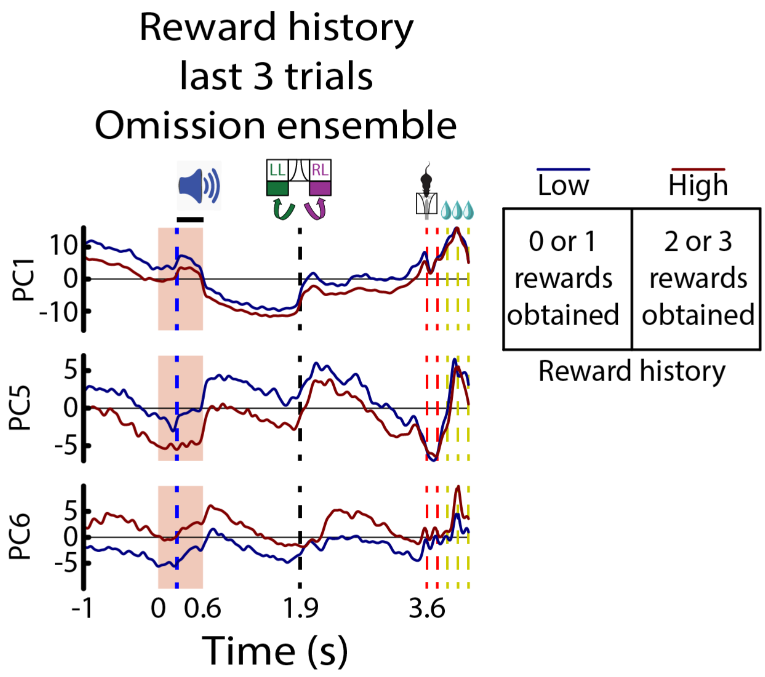


**Figure S6. The population activity of the Omission ensemble also encodes reward history.** A PCA analysis of neuronal activity within the Omission ensemble revealed distinct patterns related to recent reward history. PC1 (23.23% variance explained) and PC5 (4.87%) showed increased activity, while PC6 (4.32%) showed decreased activity when rats experienced a series of unrewarded trials. These patterns were evident throughout the trial, persisting until the first lick in the outcome epoch, suggesting the Omission ensemble encodes anticipatory signals reflecting the expectation of reward omission based on prior experience.

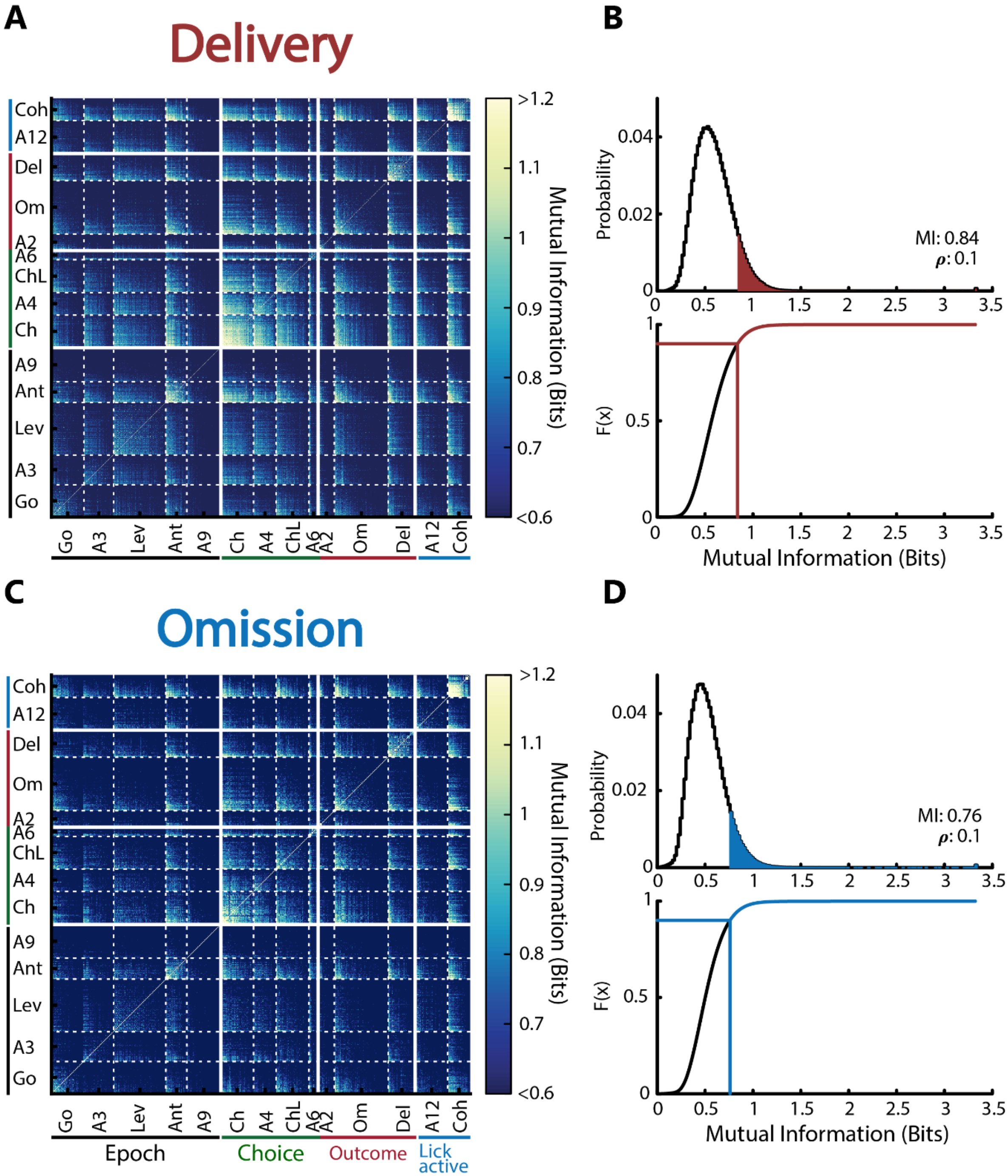


**Figure S7. Based on the Mutual Information of neuronal activity covariance in the NAcSh, we built the probabilistic network and determined the functional connectivity.**

**(A)** The heat map illustrates the mutual information between all neuron connections during reward delivery. The neuronal ensembles sort both axes in the heatmap (**Figure 4**). This metric based on mutual information between pairs of NAcSh neurons, quantifies the reduction of uncertainty or the degree of statistical dependence that one neuron provides about another. The NAcSh is a completely connected network comprising 1507 nodes and 1,134,771 connections. Thus, the functional connections are weighted and undirected.

**(B)** Top panel shows the distribution of the mutual information of all connections, whereas the bottom panel displays the cumulative density function of the same distribution. We consider it functionally "connected" if the mutual information between two neurons exceeds the 90$^{th}$ percentile of the mutual information distribution (corresponding to the blue shaded area and the blue line, top and bottom panel, respectively). Otherwise, this was considered as "disconnected."

Panels (**C**) and (**D**) have the same conventions as (**A**) and (**B**) but for reward omission.

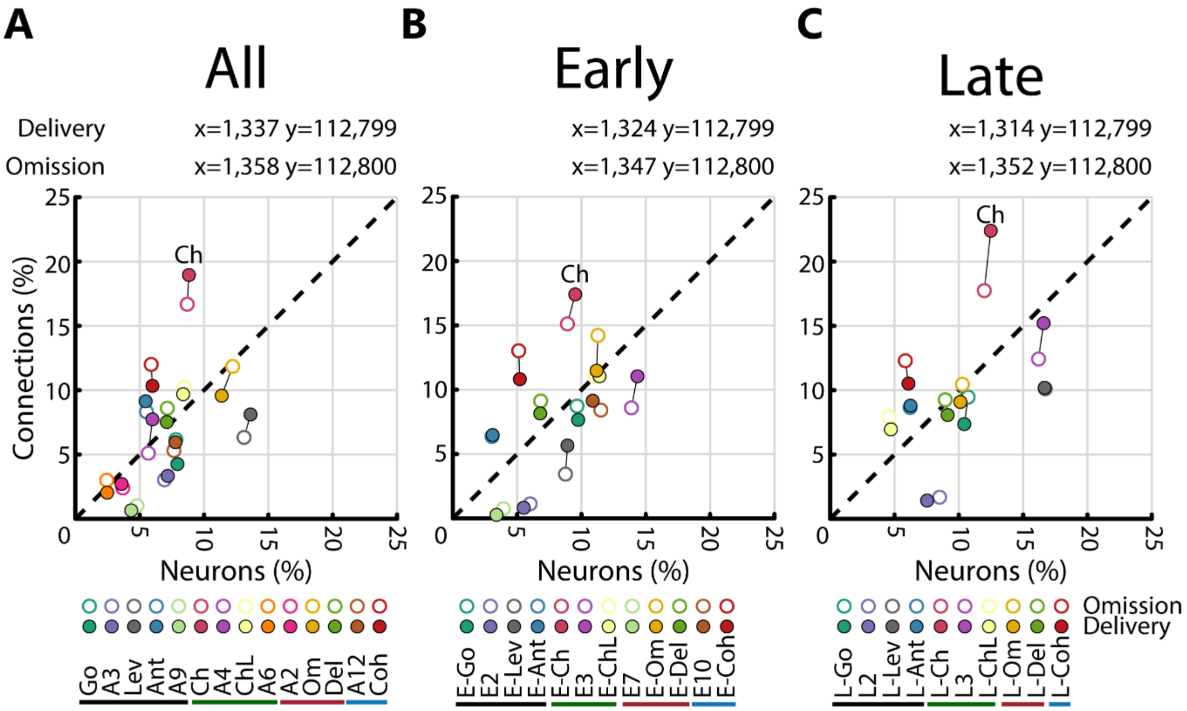


**Figure S8. The choice ensemble accounted for the largest number of connections within the NAcSh network.**

**(A-C)** Neuronal Ensembles and functional connectivity: These panels depict the relationship between the percentage of neurons and the percentage of functional connections across neuronal ensembles for All trials and during the transition from Early to Late phases. Each panel plots the number of neurons on the x-axis and connections on the y-axis, with distinct colors representing different ensembles. Filled circles represent reward delivery, while empty circles depict reward omission. The diagonal dashed line indicates equal contribution of neurons and connections.

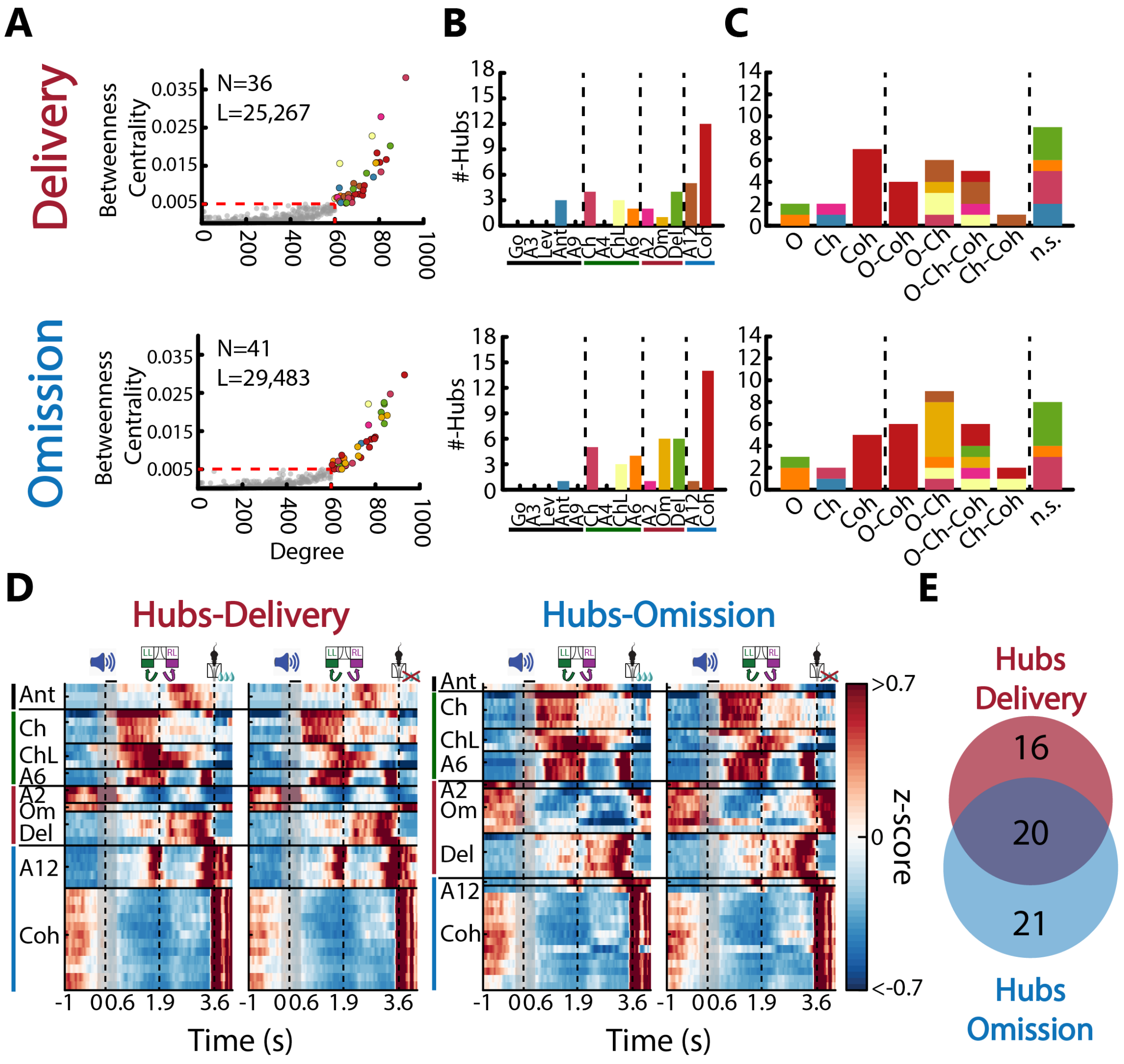


**Figure S9. Identification neuronal hubs and their neuronal responses during reward delivery and omission.**

**(A)** Neuronal hubs were identified based on two criteria: Betweenness Centrality (how often neurons lie on the shortest paths between any pair of nodes in the network; BC>0.005) and Degree connectivity (number of functional connections each neuron has; Degree>600). Neurons meeting these criteria were considered hubs and colored by ensemble membership. Otherwise, they were not considered hubs, and thus, they appeared to be colored in gray. The inset displays the number of hubs (Delivery N=36 (2.7%), Omission N=41 (3.0%)) and the number of connections these hubs have in total (Delivery L=25,267 (22.4%), Omission L=29,483 (26.14%)).

**(B)** Number of hubs across ensembles in reward delivery and omission. Bars depict the number of hubs found in each ensemble, color-coded according to the ensemble membership. Most hubs belong to lick-coherent neurons (Delivery 33.33%, Omission 34.15%).

**(C)** Behavioral variables encoded by neuronal hubs. Bars depict the number of neuronal hubs encoding reward Outcome (O), Choice (Ch), or being lick-coherent (Coh). Hubs may encode one, more than one, or none of these behavioral variables (marked by vertical dashed lines). Stacked bars are color-coded by the ensemble membership. Overall, most hubs are formed by neurons that encode reward outcome and choice (O-Ch), all behavioral variables (O-Ch-Coh), or are lick-coherent (Coh, O-Coh), and some encode none of these behavioral variables (n.s.). Additionally, the number of neurons from the Omission ensemble (Om) encoding Outcome and Choice (O-Ch) increased during the reward omission.

**(D)** Neuronal activity patterns of hubs. Left panel exhibits a heat map of activity patterns for hubs identified during reward delivery (Delivery-hubs), while the right panel exhibits hubs identified in trials where the reward was omitted (Omission-hubs). We observed that neuronal activity patterns of hubs do not necessarily vary during the outcome epoch.

**(E)** Neuronal hub overlap based on reward outcome. Only 20 neurons were hubs in both reward outcomes (20/36 Delivery, 20/41 Omission). Therefore, 57 unique hubs were identified, suggesting that different neurons propagate reward information based on the outcome.

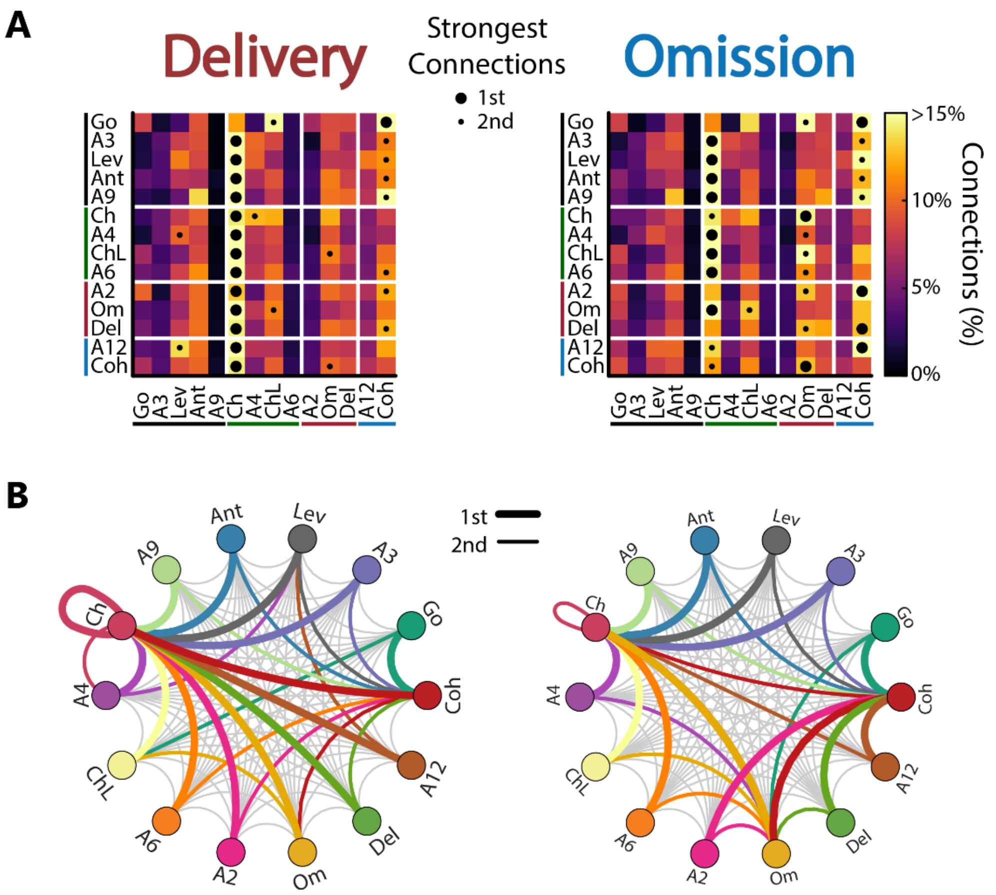


**Figure S10. The functional connectivity structure between NAcSh ensembles during reward delivery and omission**

**(A)** The heatmaps display the connectivity matrix between neuronal ensembles during reward delivery (left) and omission (right). Each cell represents the percentage of total connections accounted pairwise between neuronal ensembles. Black dots depict pairs of neuronal ensembles, with a big dot indicating the 1st strongest functional connections and a small dot indicating the 2nd strongest functional connections. In both axes, the ensembles are sorted based on meta-ensembles Epoch (black line), Choice (green line), Outcome (red line), and Lick Active (blue line). The heatmap is divided as the meta-ensembles by horizontal and vertical solid white lines.

**(B)** Bottom panels illustrate the Delivery network in a circular layout because of the ensemble's shared connections with all the remaining neuronal ensembles. The 1st and 2nd strongest functional connections are indicated by a thick (color) and thin (gray) line, respectively. The strongest connections are colored by the corresponding color assigned to each neuronal ensemble.

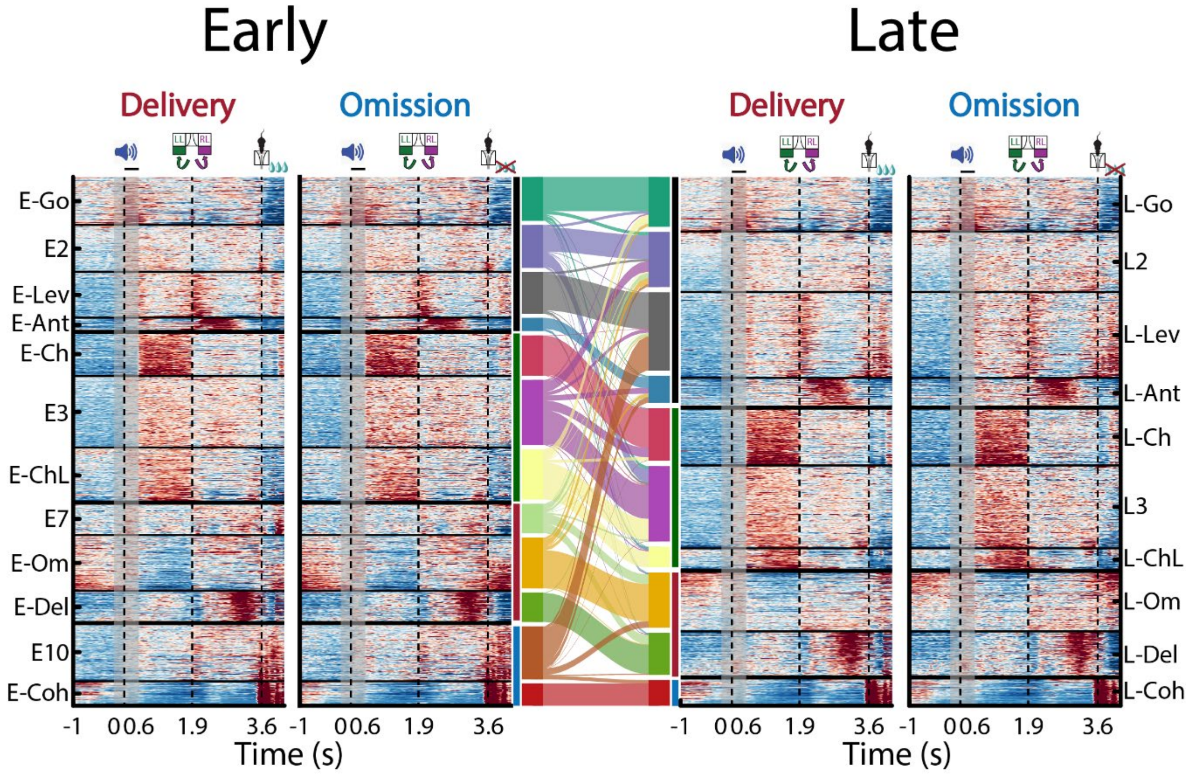


**Figure S11. Neuronal ensembles are dynamic, with neurons being removed from and added to other ensembles as reinforcement learning progresses.** Left panel, the heatmap illustrates the z-scored activity of each neuron sorted as a function of the 12 ensembles identified during the early phase. Same conventions as in **Figure 4C**. In the middle panel, the Sankey diagram draws the transition of individual neurons in each ensemble from the Early to Late phase. Right panel, the heatmap displays the z-scored activity pattern of the now 10 ensembles identified using the neural activity of the late phase. Ensembles in both task phases are sorted based on the meta-ensembles they belong to.

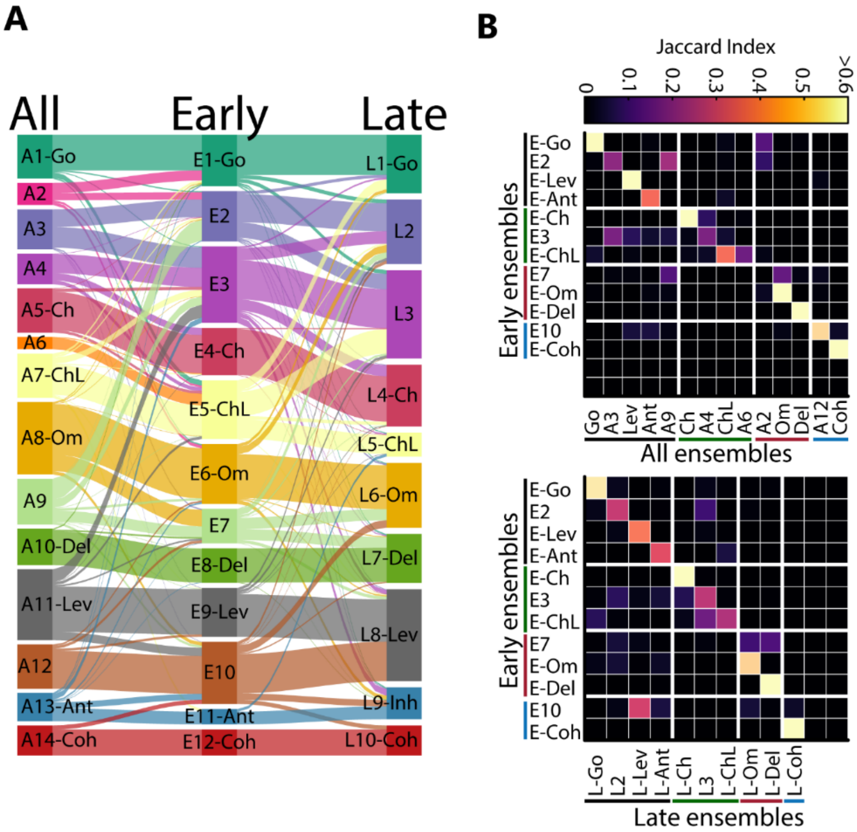


**Figure S12. Tracking individual neurons changing between neuronal ensembles found in All, Early, and Late trials of the block.**

**(A)** Sankey diagram visually represents neuronal transitions between ensembles within the block. Note the consistent coloring of ensembles across conditions.

**(B)** Comparison of neuronal ensemble overlapping in the NAcSh ensembles based on t-SNE analysis. The top panel shows the Jaccard index between the neuronal ensembles found in All trials within a block (50 trials) and those found in the Early phase (first 25 trials). The bottom panel displays the overlap between neuronal ensembles in the early and late (last 25 trials) phases. In both comparisons, the overlapping is sorted based on neighbor transitions, highlighting changes in the neuronal ensemble composition across conditions. Despite a reduction in the number of ensembles identified and by neurons moving between them, the overall transitions can be easily traced using the Jaccard index.

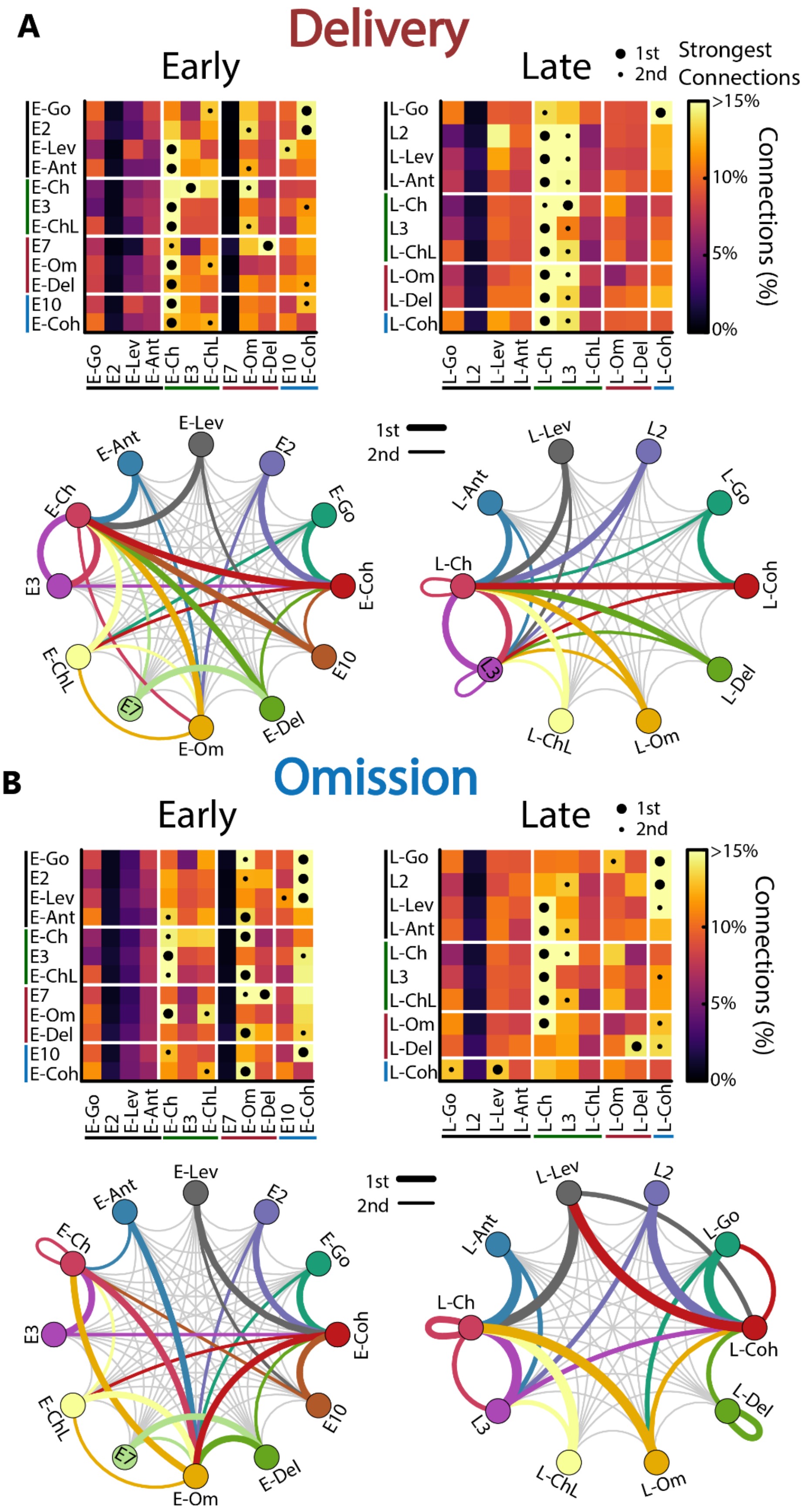
A
Delivery
Early
Late
1st
2nd
Strongest Connections
>15%
10%
5%
0%
Connections (%)
E-Go
E2
E-Lev
E-Ant
E-Ch
E3
E-ChL
E7
E-Om
E-Del
E10
E-Coh
L-Go
L2
L-Lev
L-Ant
L-Ch
L3
L-ChL
L-Om
L-Del
L-Coh
B
Omission

**Figure S13. Delivery and Omission networks during the early and late phases of reinforcement learning.**

**(A)** Delivery network reconfiguration during the transition from Early (left) to Late (right) phase. Top panels depict the connectivity matrix across ensembles and also shows the 1$^{st}$ (big dots) and 2$^{nd}$ (small dots) strongest functional connections. The connectivity matrix is divided into meta-ensembles by horizontal and vertical solid white lines. Same conventions than S10.

Bottom panels illustrate the Delivery network in a circular layout based on the ensemble's shared connections with the other neuronal ensembles. The 1$^{st}$ and 2$^{nd}$ strongest functional connections are indicated by a thick and thin line, respectively.

**(B)** Omission network reconfigures during the transition from Early (left) to Late (right) phase. Same convention as panel **A**.

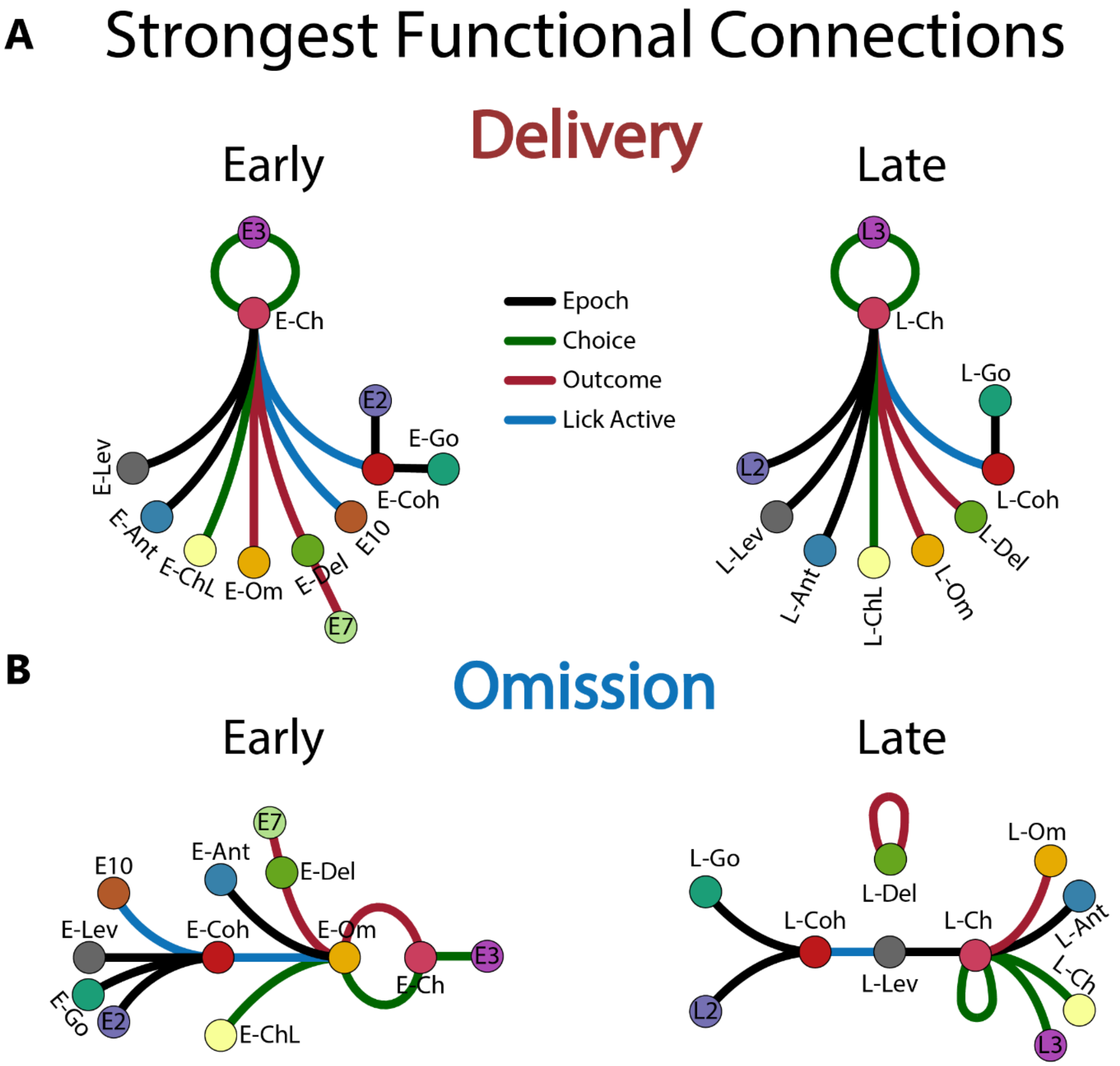


**Figure S14. Distinct neuronal ensemble structure for information flow based on reward outcome across learning.**

**(A)** The NAcSh network reshapes its connectivity during reward delivery. Circles are color-coded according to the ensemble membership. Thick lines depict the strongest functional connections, with line colors indicating the Meta-Ensemble to which it belongs. The network structure during reward delivery representation in the NAcSh exhibited how the modular reward organization revolves around the Choice ensemble (E-Ch and L-Ch, left and right panel, respectively) and forms a close loop with neuronal ensemble 3 (E3 and L3).

**(B)** Functional connectivity reorganization in the Omission network. Same conventions as in panel (**A**). Unlike reward delivery, the network structure during reward omission representation in the NAcSh undergoes changes through learning. In the early phase, neuronal ensembles appear organized hierarchically, with significant functional connections between E-Om and E-Ch. In the late phase, core connections resemble an interface between L-Ch with L-Coh, facilitating communication between decision variables and licking behavior. Additionally, the L-Del ensemble emerges as an isolated module during reward omission.
