## Supplementary material for "The flow of reward information through neuronal ensembles in the accumbens": Document S1

| *Network* | ***Power-law fit*** | ***Power-law*** | ***Log-Normal*** | ***Poisson*** |
|---|---|---|---|---|
| | **γ** | | *Probability* | |
| All-Average | 0.7277 | **0.9252** | 0.1608 | 0.2796 |
| All-Delivery | 0.6931 | 0.2386 | 0.0116 | **0.3084** |
| All-Omission | 0.7041 | 0.1482 | 0.0026 | **0.2082** |
| Early-Average | 0.6877 | 0.2628 | 0.024 | **0.5074** |
| Early-Delivery | 0.7024 | 0.096 | 0.0134 | **0.5962** |
| Early-Omission | 0.6877 | 0 | **0.0108** | 0 |
| Late-Average | 0.6728 | 0.0078 | **0.806** | 0.658 |
| Late-Delivery | 0.6731 | 0.0668 | 0.0142 | **0.4702** |
| Late-Omission | 0.7209 | 0.0154 | **0.8298** | 0.5092 |

#### Power law long-tailed degree distributions

In the context of a network's degree distribution, if the tail follows a power-law distribution, it indicates that there are a few nodes (hubs) with extremely high degrees, and the frequency of nodes with high degrees decreases according to a power-law relationship. In other words, a small number of nodes have a disproportionately large number of connections. This observation is often associated with scale-free networks, which are characterized by the presence of hubs that play a critical role in the network's structure and function. The power-law behavior in the degree distribution suggests that there are no characteristic scales in the network, and the connectivity patterns are self-similar across different scales.

#### Poisson long-tailed degree distributions

A Poisson distribution is a probability distribution that expresses the number of events that occur within a fixed interval of time or space. It is often used to model rare events that occur independently. In the context of a network, if the high-degree nodes (nodes with many connections) exhibit a distribution pattern similar to a Poisson distribution, it suggests that the occurrence of nodes with extremely high degrees is relatively rare and follows a certain probabilistic pattern.

In simpler terms, it indicates that most nodes in the network have a moderate number of connections, but as you move to nodes with very high degrees (the tail of the distribution), the probability of finding nodes with an exceptionally high degree decreases according to a Poisson-like pattern. This kind of behavior is often observed in some real-world networks, where the majority of nodes have only a few connections, but there are a few nodes (hubs) with significantly higher degrees.

#### Log-normal long-tailed degree distributions

A lognormal distribution is a probability distribution that is associated with a random variable whose logarithm is normally distributed. In the context of a network, if the degrees of high-degree nodes (nodes with many connections) follow a lognormal distribution in their statistical behavior, it implies that the logarithms of these degrees are normally distributed. In practical terms, this suggests that the high-degree nodes in the network have a multiplicative effect on their degrees, and the distribution of their degrees follows a pattern that resembles a bell-shaped curve when plotted on a logarithmic scale. If the tail of the degree distribution follows a lognormal distribution, it implies that the extremely high-degree nodes in the network follow a statistical pattern where the logarithms of their degrees have a normal distribution. This behavior is characteristic of certain complex networks, particularly those with scale-free properties.

### PowerLaws_Neuro.R

enrique

2024-02-06

```r
# Power law distribution empirical verification using the poweRlaw R library
# https://github.com/csgillespie/poweRlaw
# Based on the work of Clauset, Shalizi, and Newman
# https://aaronclauset.github.io/powerlaws/

#install.packages("poweRlaw")
library("poweRlaw")
setwd("/Users/enrique/Dropbox/Ranier_modelos/PowerLawDistributions")

#################################################################
# Actual networks from the paper "The flow of pleasure through  ##
# neuronal ensembles in the accumbens" ##########################
#################################################################

# Average all

allavg <-read.csv(file="network_stats/Node_Average_all.csv", header = TRUE)
#str(allavg)

#Select and check the degree column
degavg <- allavg$Degree
degavg <-sort(degavg)
head(degavg)
```

```
## [1] 1 1 1 1 1 1
```

```r
tail(degavg)
```

```
## [1] 806 821 841 899 911 961
```

```r
# Remove entries equal to zero

#degavg <-degavg[degavg!=0]

#Fit a maximum likelihood discrete powerlaw distribution form the data

m_pl=displ$new(degavg)

# Infer model parameters (lower threshold)

est=estimate_xmin(m_pl)

# Update the powerlaw distribution minimum value
```

```
m_pl$setXmin(est)

# Fit alternative distributions, e.g. a lognormal and a Poison

m_ln=dislnorm$new(degavg)
est=estimate_xmin(m_ln)
m_ln$setXmin(est)

m_pois=dispois$new(degavg)
est=estimate_xmin(m_pois)
m_pois$setXmin(est)


# Plot the competing distributions

plot(m_pl)
lines(m_pl,col=2)
lines(m_ln,col=3)
lines(m_pois,col=4)
```

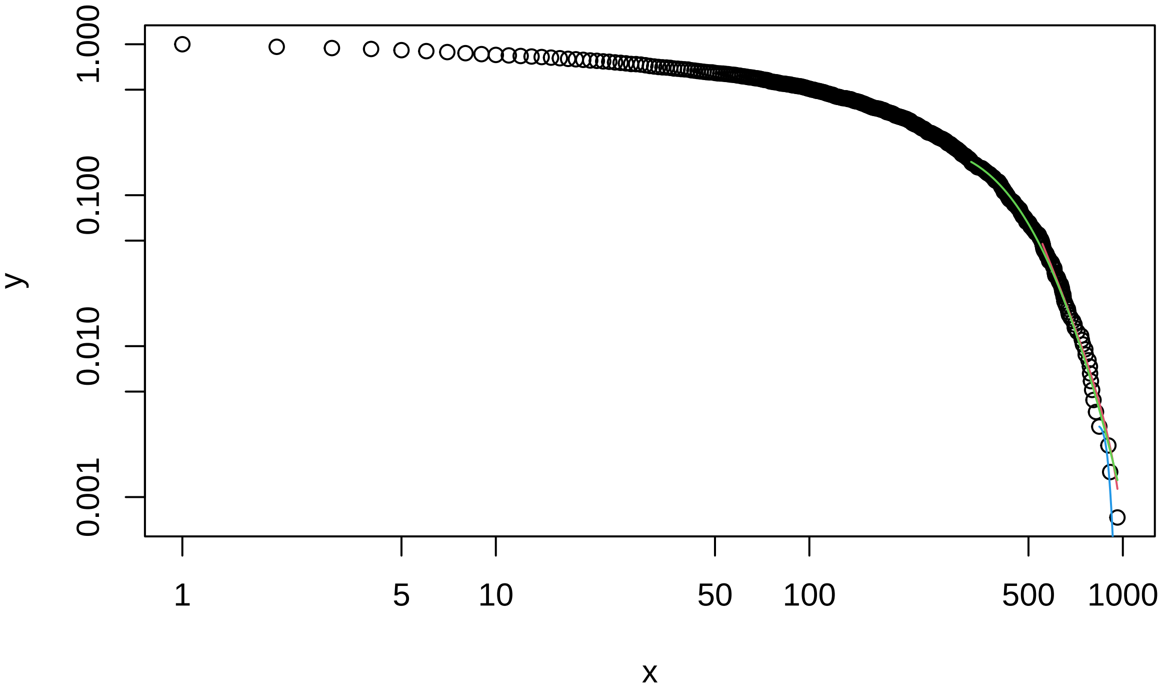


```
# Testing the Powerlaw hypothesis using a bootstrap function

bs_p=bootstrap_p(m_pl, no_of_sims =5000, threads=10)
```

```
## Expected total run time for 5000 sims, using 10 threads is 3300 seconds.
```

```
bs_p$p
```

```
## [1] 0.9252
```

```
# Testing the other distributions P is the a posteriori probability that the
#data comes from this distribution
```

```
#LogNormal
bs_ln=bootstrap_p(m_ln,  no_of_sims =5000, threads=10)
```

```
## Expected total run time for 5000 sims, using 10 threads is 40200 seconds.
```

```
bs_ln$p
```

```
## [1] 0.1608
```

```
# Poison
bs_pois=bootstrap_p(m_pois, no_of_sims =5000,  threads=10)
```

```
## Expected total run time for 5000 sims, using 10 threads is 1000 seconds.
```

```
bs_pois$p
```

```
## [1] 0.2796
```

```
# Average early

earlyavg <-read.csv(file="network_stats/Node_Average_early.csv", header = TRUE)
#str(earlyavg)

#Select and check the degree column
degear <- earlyavg$Degree
degear <-sort(degear)
head(degear)
```

```
## [1] 1 1 1 1 1 1
```

```
tail(degear)
```

```
## [1] 823 837 839 859 865 872
```

```
# Remove entries equal to zero

#degear <-degear[degear!=0]

#Fit a maximum likelihood discrete powerlaw distribution form the data

m_pl=displ$new(degear)

# Infer model parameters (lower threshold)

est=estimate_xmin(m_pl)

# Update the powerlaw distribution minimum value

m_pl$setXmin(est)

# Fit alternative distributions, e.g. a lognormal and a Poison

m_ln=dislnorm$new(degear)
est=estimate_xmin(m_ln)
m_ln$setXmin(est)

m_pois=dispois$new(degear)
est=estimate_xmin(m_pois)
```

```r
m_pois$setXmin(est)


# Plot the competing distributions

plot(m_pl)
lines(m_pl,col=2)
lines(m_ln,col=3)
lines(m_pois,col=4)
```

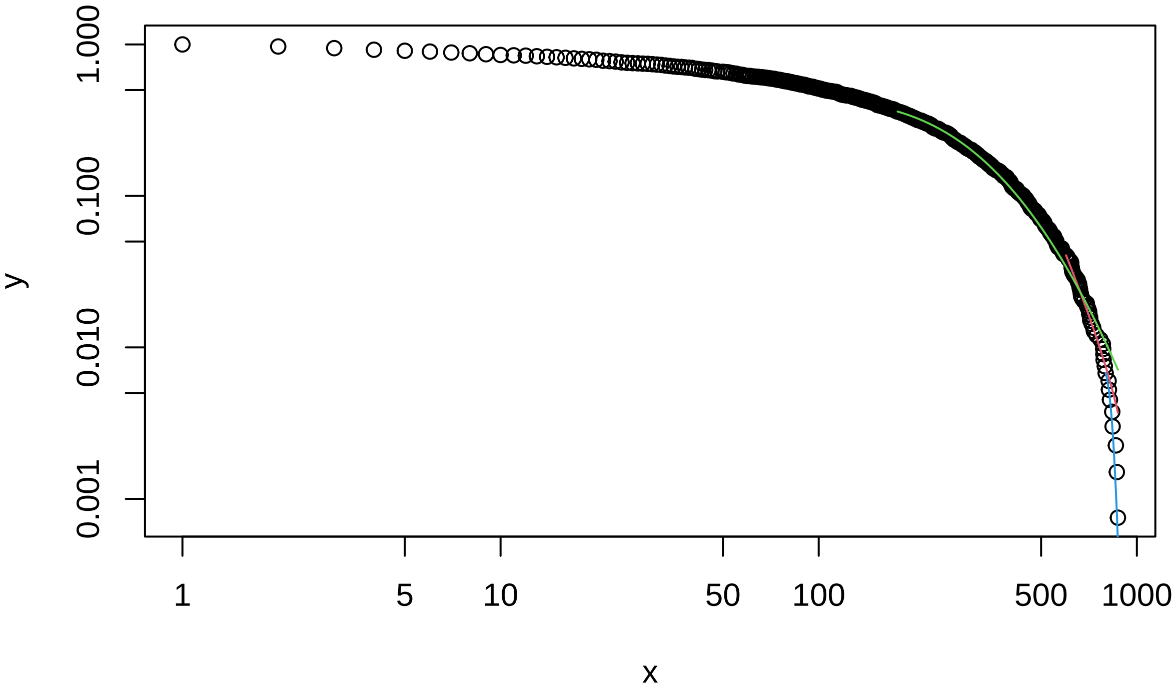


```r
# Testing the  Powerlaw hypothesis using a bootstrap function

bs_p=bootstrap_p(m_pl, no_of_sims =5000, threads=10)
```

```
## Expected total run time for 5000 sims, using 10 threads is 3510 seconds.
```

```r
bs_p$p
```

```
## [1] 0.2628
```

```r
# Testing the other distributions P is the a posteriori probability that the
#data comes from this distribution

#LogNormal
bs_ln=bootstrap_p(m_ln, no_of_sims =5000, threads=10)
```

```
## Expected total run time for 5000 sims, using 10 threads is 6770 seconds.
```

```r
bs_ln$p
```

```
## [1] 0.024
```

```r
# Poison
bs_pois=bootstrap_p(m_pois, no_of_sims =5000, threads=10)
```

```
## Expected total run time for 5000 sims, using 10 threads is 1020 seconds.
```

```
bs_pois$p
```

```
## [1] 0.5074
```

```
# Average late

lateavg <-read.csv(file="network_stats/Node_Average_late.csv", header = TRUE)
#str(lateavg)

#Select and check the degree column
deglat <- lateavg$Degree
deglat <-sort(deglat)
head(deglat)
```

```
## [1] 1 1 1 1 1 1
```

```
tail(deglat)
```

```
## [1] 828 845 848 851 866 888
```

```
# Remove entries equal to zero

#deglat <-deglat[deglat!=0]

#Fit a maximum likelihood discrete powerlaw distribution form the data

m_pl=displ$new(deglat)

# Infer model parameters (lower threshold)

est=estimate_xmin(m_pl)

# Update the powerlaw distribution minimum value

m_pl$setXmin(est)

# Fit alternative distributions, e.g. a lognormal and a Poison

m_ln=dislnorm$new(deglat)
est=estimate_xmin(m_ln)
m_ln$setXmin(est)

m_pois=dispois$new(deglat)
est=estimate_xmin(m_pois)
m_pois$setXmin(est)


# Plot the competing distributions

plot(m_pl)
lines(m_pl,col=2)
lines(m_ln,col=3)
lines(m_pois,col=4)
```

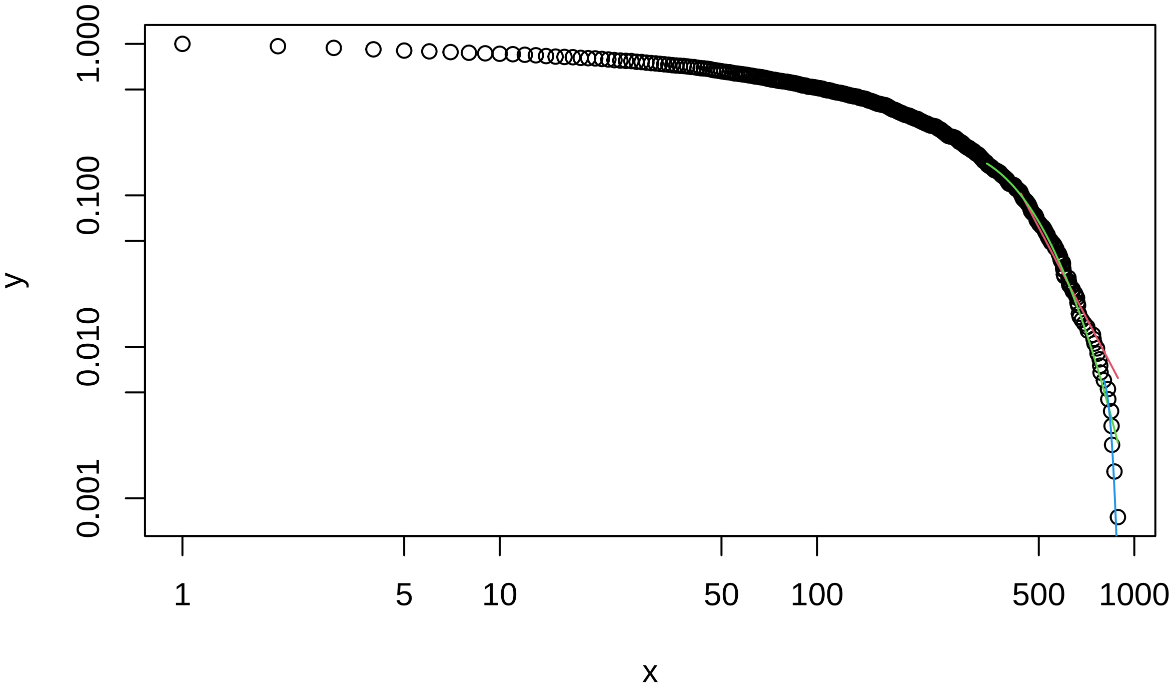


```r
# Testing the  Powerlaw hypothesis using a bootstrap function

bs_p=bootstrap_p(m_pl, no_of_sims =5000, threads=10)
```

```
## Expected total run time for 5000 sims, using 10 threads is 3370 seconds.
```

```r
bs_p$p
```

```
## [1] 0.0078
```

```r
# Testing the other distributions P is the a posteriori probability that the
#data comes from this distribution

#LogNormal
bs_ln=bootstrap_p(m_ln, no_of_sims =5000, threads=10)
```

```
## Expected total run time for 5000 sims, using 10 threads is 6260 seconds.
```

```r
bs_ln$p
```

```
## [1] 0.806
```

```r
# Poison
bs_pois=bootstrap_p(m_pois, no_of_sims =5000, threads=10)
```

```
## Expected total run time for 5000 sims, using 10 threads is 1130 seconds.
```

```r
bs_pois$p
```

```
## [1] 0.658
```

```r
# Delivery all

delall <-read.csv(file="network_stats/Node_Delivery_all.csv", header = TRUE)
#str(delall)

#Select and check the degree column
```

```r
degdelall <- delall$Degree
degdelall <-sort(degdelall)
head(degdelall)
```

```
## [1] 1 1 1 1 1 1
```

```r
tail(degdelall)
```

```
## [1] 803 808 808 832 850 920
```

```r
# Remove entries equal to zero

#degdelall <-degdelall[degdelall!=0]

#Fit a maximum likelihood discrete powerlaw distribution form the data

m_pl=displ$new(degdelall)

# Infer model parameters (lower threshold)

est=estimate_xmin(m_pl)

# Update the powerlaw distribution minimum value

m_pl$setXmin(est)

# Fit alternative distributions, e.g. a lognormal and a Poison

m_ln=dislnorm$new(degdelall)
est=estimate_xmin(m_ln)
m_ln$setXmin(est)

m_pois=dispois$new(degdelall)
est=estimate_xmin(m_pois)
m_pois$setXmin(est)


# Plot the competing distributions

plot(m_pl)
lines(m_pl,col=2)
lines(m_ln,col=3)
lines(m_pois,col=4)
```

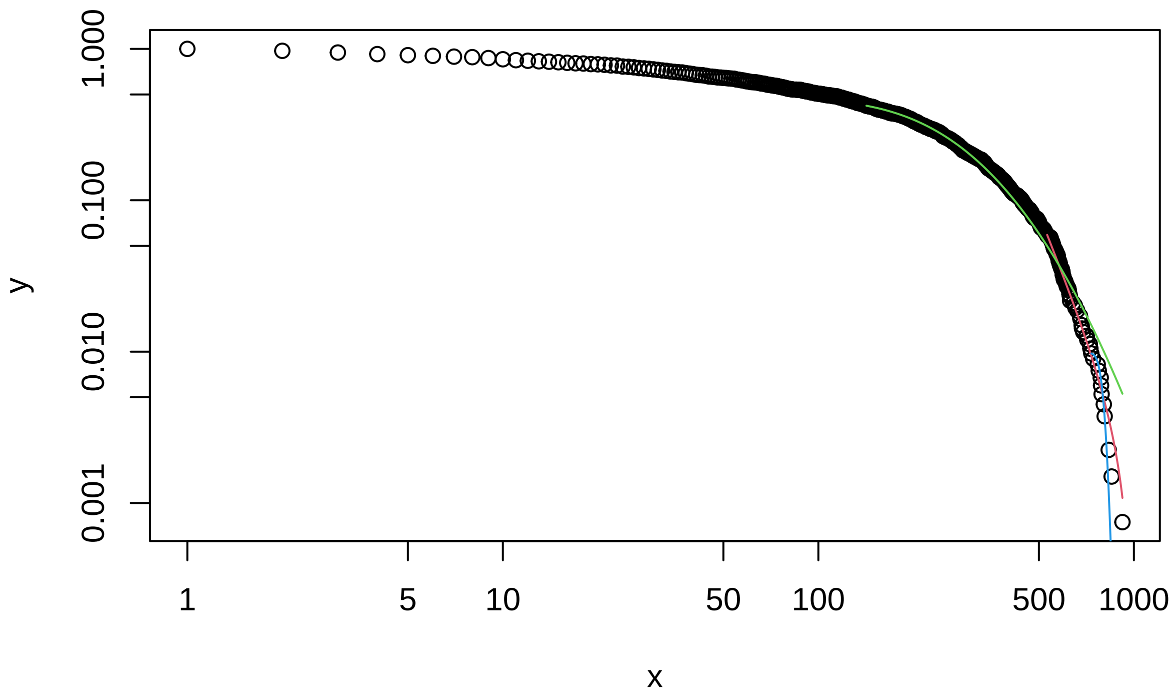


```
# Testing the  Powerlaw hypothesis using a bootstrap function

bs_p=bootstrap_p(m_pl, no_of_sims =5000, threads=10)
```

```
## Expected total run time for 5000 sims, using 10 threads is 3600 seconds.
```

```
bs_p$p
```

```
## [1] 0.2386
```

```
# Testing the other distributions P is the a posteriori probability that the
#data comes from this distribution

#LogNormal
bs_ln=bootstrap_p(m_ln, no_of_sims =5000, threads=10)
```

```
## Expected total run time for 5000 sims, using 10 threads is 16900 seconds.
```

```
bs_ln$p
```

```
## [1] 0.0116
```

```
# Poison
bs_pois=bootstrap_p(m_pois, no_of_sims =5000, threads=10)
```

```
## Expected total run time for 5000 sims, using 10 threads is 973 seconds.
```

```
bs_pois$p
```

```
## [1] 0.3084
```

```
# Delivery early

delear <-read.csv(file="network_stats/Node_Delivery_early.csv", header = TRUE)
#str(delear)

#Select and check the degree column
```

```r
degdelear <- delear$Degree
degdelear <-sort(degdelear)
head(degdelear)
```

```
## [1] 1 1 1 1 1 1
```

```r
tail(degdelear)
```

```
## [1] 819 831 854 865 882 914
```

```r
# Remove entries equal to zero

#degdelear <-degdelear[degdelear!=0]

#Fit a maximum likelihood discrete powerlaw distribution form the data

m_pl=displ$new(degdelear)

# Infer model parameters (lower threshold)

est=estimate_xmin(m_pl)

# Update the powerlaw distribution minimum value

m_pl$setXmin(est)

# Fit alternative distributions, e.g. a lognormal and a Poison

m_ln=dislnorm$new(degdelear)
est=estimate_xmin(m_ln)
m_ln$setXmin(est)

m_pois=dispois$new(degdelear)
est=estimate_xmin(m_pois)
m_pois$setXmin(est)


# Plot the competing distributions

plot(m_pl)
lines(m_pl,col=2)
lines(m_ln,col=3)
lines(m_pois,col=4)
```

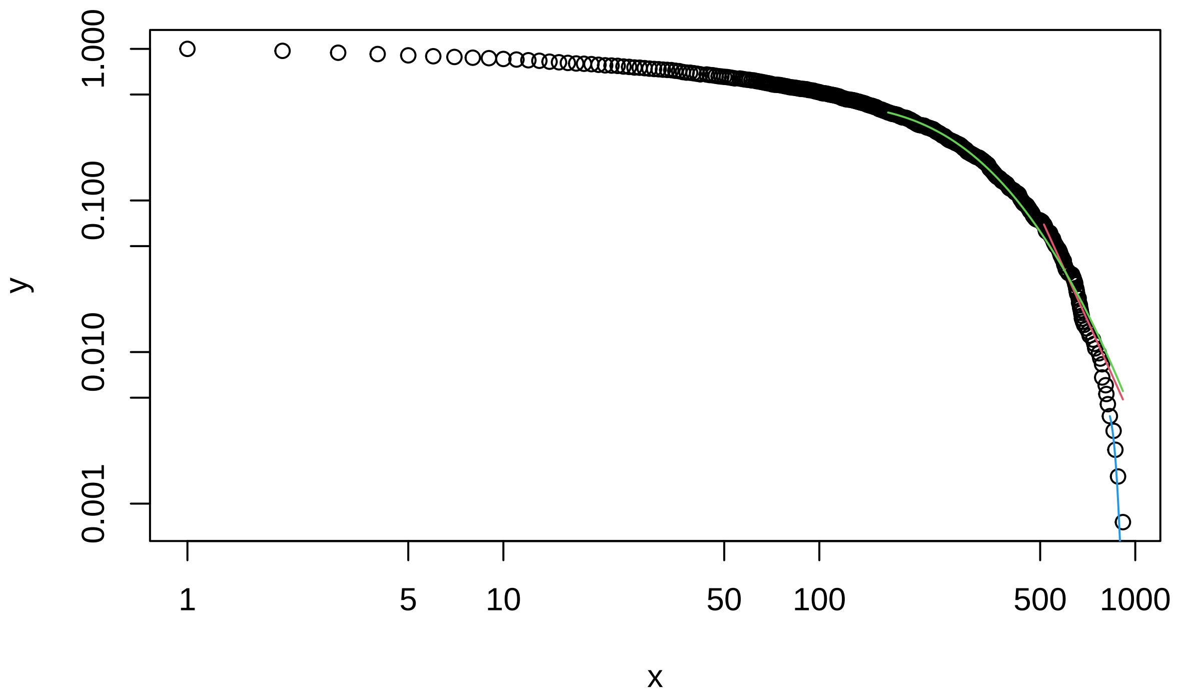


```
# Testing the  Powerlaw hypothesis using a bootstrap function

bs_p=bootstrap_p(m_pl, no_of_sims =5000, threads=10)
```

```
## Expected total run time for 5000 sims, using 10 threads is 3190 seconds.
```

```
bs_p$p
```

```
## [1] 0.096
```

```
# Testing the other distributions P is the a posteriori probability that the
#data comes from this distribution

#LogNormal
bs_ln=bootstrap_p(m_ln, no_of_sims =5000, threads=10)
```

```
## Expected total run time for 5000 sims, using 10 threads is 6350 seconds.
```

```
bs_ln$p
```

```
## [1] 0.0134
```

```
# Poison
bs_pois=bootstrap_p(m_pois, no_of_sims =5000, threads=10)
```

```
## Expected total run time for 5000 sims, using 10 threads is 946 seconds.
```

```
bs_pois$p
```

```
## [1] 0.5962
```

```
# Delivery late


delate <-read.csv(file="network_stats/Node_Delivery_late.csv", header = TRUE)
#str(delate)
```

```r
#Select and check the degree column
degdelate <- delate$Degree
degdelate <-sort(degdelate)
head(degdelate)
```

```
## [1] 1 1 1 1 1 1
```

```r
tail(degdelate)
```

```
## [1] 798 800 803 809 833 860
```

```r
# Remove entries equal to zero

#degdelate <-degdelate[degdelate!=0]

#Fit a maximum likelihood discrete powerlaw distribution form the data

m_pl=displ$new(degdelate)

# Infer model parameters (lower threshold)

est=estimate_xmin(m_pl)

# Update the powerlaw distribution minimum value

m_pl$setXmin(est)

# Fit alternative distributions, e.g. a lognormal and a Poison

m_ln=dislnorm$new(degdelate)
est=estimate_xmin(m_ln)
m_ln$setXmin(est)

m_pois=dispois$new(degdelate)
est=estimate_xmin(m_pois)
m_pois$setXmin(est)


# Plot the competing distributions

plot(m_pl)
lines(m_pl,col=2)
lines(m_ln,col=3)
lines(m_pois,col=4)
```

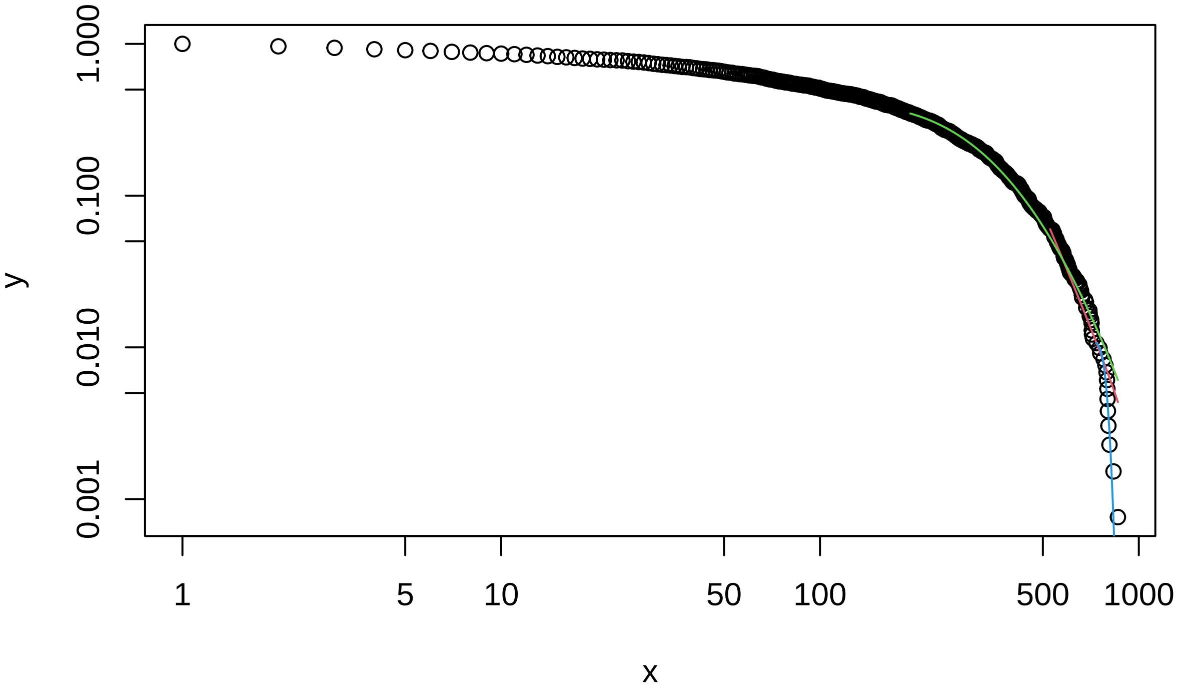


```
# Testing the  Powerlaw hypothesis using a bootstrap function

bs_p=bootstrap_p(m_pl, no_of_sims =5000, threads=10)
```

```
## Expected total run time for 5000 sims, using 10 threads is 3500 seconds.
```

```
bs_p$p
```

```
## [1] 0.0668
```

```
# Testing the other distributions P is the a posteriori probability that the
#data comes from this distribution

#LogNormal
bs_ln=bootstrap_p(m_ln, no_of_sims =5000, threads=10)
```

```
## Expected total run time for 5000 sims, using 10 threads is 6800 seconds.
```

```
bs_ln$p
```

```
## [1] 0.0142
```

```
# Poison
bs_pois=bootstrap_p(m_pois, no_of_sims =5000, threads=10)
```

```
## Expected total run time for 5000 sims, using 10 threads is 1130 seconds.
```

```
bs_pois$p
```

```
## [1] 0.4702
```

```
# Omission all

omall <-read.csv(file="network_stats/Node_Omission_all.csv", header = TRUE)
#str(delate)

#Select and check the degree column
```

```r
degomall <- omall$Degree
degomall <-sort(degomall)
head(degomall)
```

```
## [1] 1 1 1 1 1 1
```

```r
tail(degomall)
```

```
## [1] 839 840 840 853 867 931
```

```r
# Remove entries equal to zero

#degdelate <-degdelate[degdelate!=0]

#Fit a maximum likelihood discrete powerlaw distribution form the data

m_pl=displ$new(degomall)

# Infer model parameters (lower threshold)

est=estimate_xmin(m_pl)

# Update the powerlaw distribution minimum value

m_pl$setXmin(est)

# Fit alternative distributions, e.g. a lognormal and a Poison

m_ln=dislnorm$new(degomall)
est=estimate_xmin(m_ln)
m_ln$setXmin(est)

m_pois=dispois$new(degomall)
est=estimate_xmin(m_pois)
m_pois$setXmin(est)


# Plot the competing distributions

plot(m_pl)
lines(m_pl,col=2)
lines(m_ln,col=3)
lines(m_pois,col=4)
```

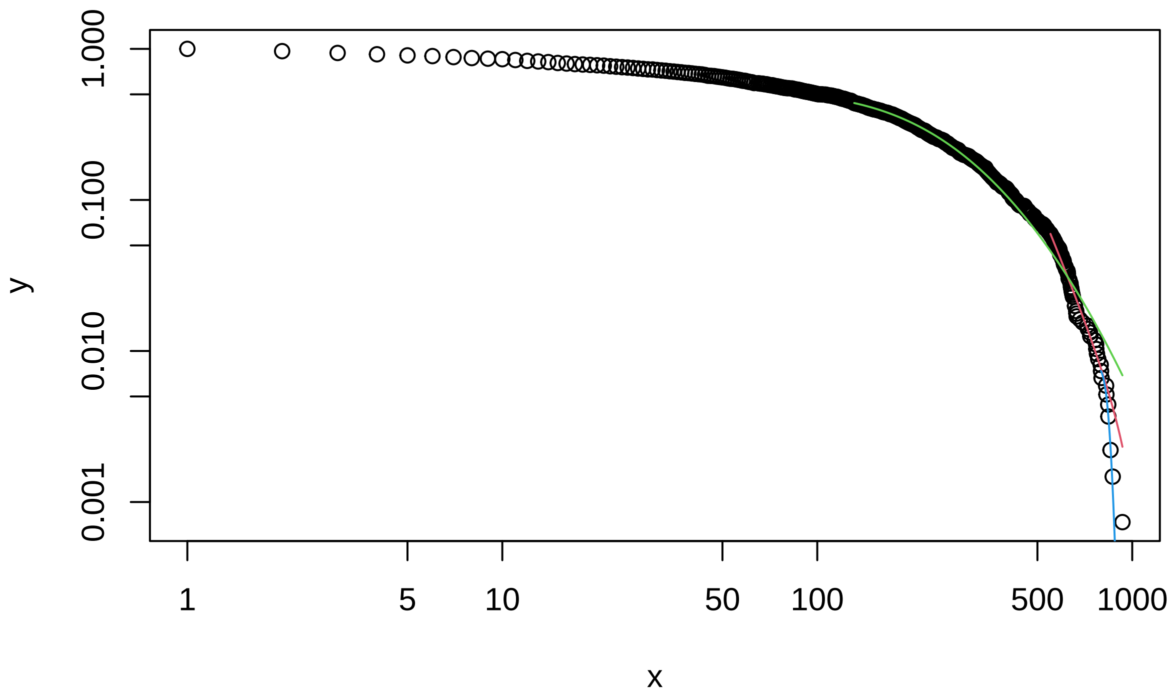


```
# Testing the  Powerlaw hypothesis using a bootstrap function

bs_p=bootstrap_p(m_pl, no_of_sims =5000, threads=10)
```

```
## Expected total run time for 5000 sims, using 10 threads is 3000 seconds.
```

```
bs_p$p
```

```
## [1] 0.1482
```

```
# Testing the other distributions P is the a posteriori probability that the
#data comes from this distribution

#LogNormal
bs_ln=bootstrap_p(m_ln, no_of_sims =5000, threads=10)
```

```
## Expected total run time for 5000 sims, using 10 threads is 6060 seconds.
```

```
bs_ln$p
```

```
## [1] 0.0026
```

```
# Poison
bs_pois=bootstrap_p(m_pois, no_of_sims =5000, threads=10)
```

```
## Expected total run time for 5000 sims, using 10 threads is 940 seconds.
```

```
bs_pois$p
```

```
## [1] 0.2082
```

```
# Omission early

omear <-read.csv(file="network_stats/Node_Omission_early.csv", header = TRUE)
#str(delate)

#Select and check the degree column
```

```r
degomear <- omear$Degree
degomear <-sort(degomear)
head(degomear)
```

```
## [1] 1 1 1 1 1 1
```

```r
tail(degomear)
```

```
## [1] 814 823 849 875 922 932
```

```r
# Remove entries equal to zero

#degdelate <-degdelate[degdelate!=0]

#Fit a maximum likelihood discrete powerlaw distribution form the data

m_pl=displ$new(degomear)

# Infer model parameters (lower threshold)

est=estimate_xmin(m_pl)

# Update the powerlaw distribution minimum value

m_pl$setXmin(est)

# Fit alternative distributions, e.g. a lognormal and a Poison

m_ln=dislnorm$new(degomear)
est=estimate_xmin(m_ln)
m_ln$setXmin(est)

m_pois=dispois$new(degomear)
est=estimate_xmin(m_pois)
m_pois$setXmin(est)


# Plot the competing distributions

plot(m_pl)
lines(m_pl,col=2)
lines(m_ln,col=3)
lines(m_pois,col=4)
```

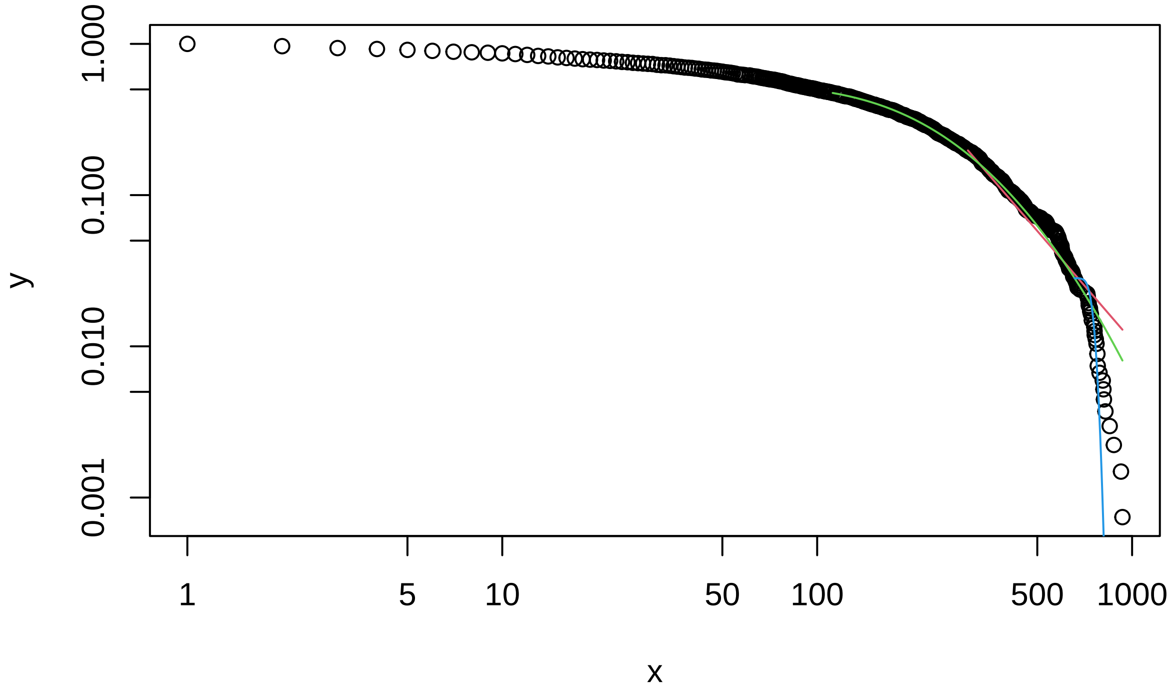


```
# Testing the  Powerlaw hypothesis using a bootstrap function

bs_p=bootstrap_p(m_pl, no_of_sims =5000, threads=10)
```

```
## Expected total run time for 5000 sims, using 10 threads is 2910 seconds.
```

```
bs_p$p
```

```
## [1] 0
```

```
# Testing the other distributions P is the a posteriori probability that the
#data comes from this distribution

#LogNormal
bs_ln=bootstrap_p(m_ln, no_of_sims =5000, threads=10)
```

```
## Expected total run time for 5000 sims, using 10 threads is 6060 seconds.
```

```
bs_ln$p
```

```
## [1] 0.0108
```

```
# Poison
bs_pois=bootstrap_p(m_pois, no_of_sims =5000, threads=10)
```

```
## Expected total run time for 5000 sims, using 10 threads is 979 seconds.
```

```
bs_pois$p
```

```
## [1] 0
```

```
# Omission late

omlate <-read.csv(file="network_stats/Node_Omission_late.csv", header = TRUE)
#str(delate)

#Select and check the degree column
```

```r
degomlate <- omlate$Degree
degomlate <-sort(degomlate)
head(degomlate)
```

```
## [1] 1 1 1 1 1 1
```

```r
tail(degomlate)
```

```
## [1] 845 856 859 861 865 944
```

```r
# Remove entries equal to zero

#degdelate <-degdelate[degdelate!=0]

#Fit a maximum likelihood discrete powerlaw distribution form the data

m_pl=displ$new(degomlate)

# Infer model parameters (lower threshold)

est=estimate_xmin(m_pl)

# Update the powerlaw distribution minimum value

m_pl$setXmin(est)

# Fit alternative distributions, e.g. a lognormal and a Poison

m_ln=dislnorm$new(degomlate)
est=estimate_xmin(m_ln)
m_ln$setXmin(est)

m_pois=dispois$new(degomlate)
est=estimate_xmin(m_pois)
m_pois$setXmin(est)


# Plot the competing distributions

plot(m_pl)
lines(m_pl,col=2)
lines(m_ln,col=3)
lines(m_pois,col=4)
```

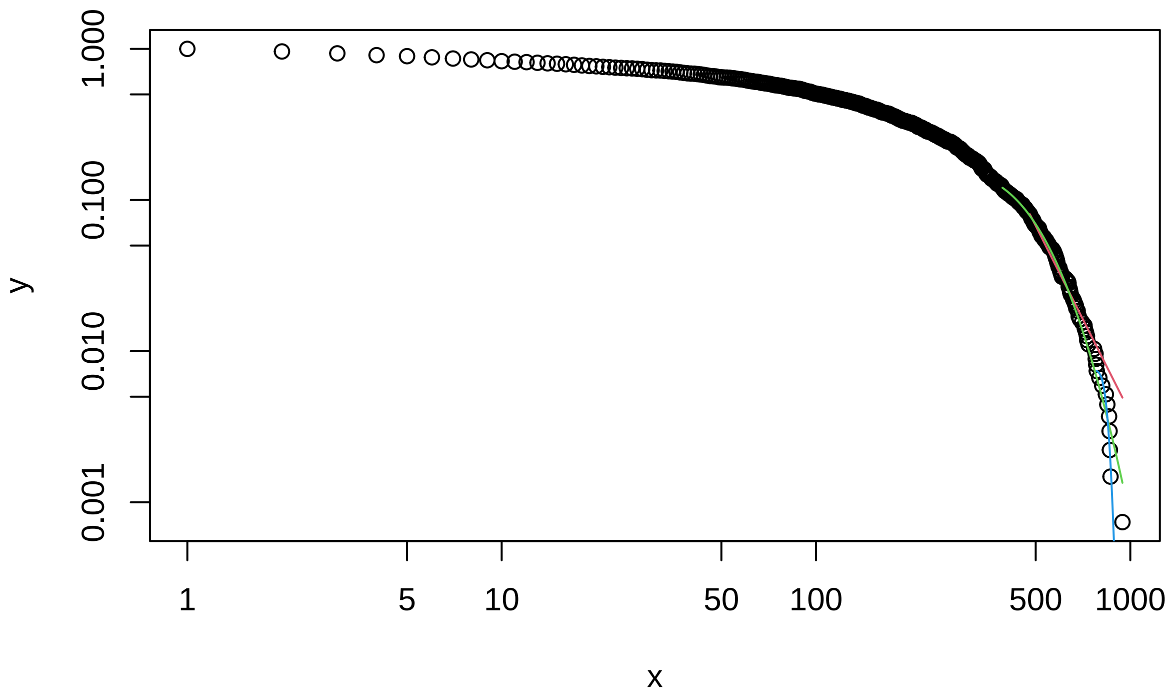


```
# Testing the  Powerlaw hypothesis using a bootstrap function

bs_p=bootstrap_p(m_pl, no_of_sims =5000, threads=10)
```

```
## Expected total run time for 5000 sims, using 10 threads is 3140 seconds.
```

```
bs_p$p
```

```
## [1] 0.0154
```

```
# Testing the other distributions P is the a posteriori probability that the
#data comes from this distribution

#LogNormal
bs_ln=bootstrap_p(m_ln, no_of_sims =5000, threads=10)
```

```
## Expected total run time for 5000 sims, using 10 threads is 6020 seconds.
```

```
bs_ln$p
```

```
## [1] 0.8298
```

```
# Poison
bs_pois=bootstrap_p(m_pois, no_of_sims =5000, threads=10)
```

```
## Expected total run time for 5000 sims, using 10 threads is 963 seconds.
```

```
bs_pois$p
```

```
## [1] 0.5092
```
